# Mutations of the YidC Insertase alleviate stress from σ^M^-dependent membrane protein overproduction in *Bacillus subtilis*

**DOI:** 10.1101/678235

**Authors:** Heng Zhao, Ankita J. Sachla, John D. Helmann

## Abstract

In *Bacillus subtilis*, the extracytoplasmic function σ factor σ^M^ regulates cell wall synthesis and is critical for intrinsic resistance to cell wall targeting antibiotics. The anti-σ factors YhdL and YhdK form a complex that restricts the basal activity of σ^M^, and the absence of YhdL leads to runaway expression of the σ^M^ regulon and cell death. Here, we report that this lethality can be suppressed by gain-of-function mutations in *spoIIIJ*, which encodes the major YidC membrane protein insertase in *B. subtilis*. *B. subtilis* PY79 SpoIIIJ contains a single amino acid substitution in the substrate-binding channel (Q140K), and this allele suppresses the lethality of high SigM. Analysis of a library of YidC variants reveals that increased charge (+2 or +3) in the substrate-binding channel can compensate for high expression of the σ^M^ regulon. Derepression of the σ^M^ regulon induces secretion stress, oxidative stress and DNA damage responses, all of which can be alleviated by the YidC^Q140K^ substitution. We further show that the fitness defect caused by high σ^M^ activity is exacerbated in the absence of SecDF protein translocase or σ^M^-dependent induction of the Spx oxidative stress regulon. Conversely, cell growth is improved by mutation of specific σ^M^-dependent promoters controlling operons encoding integral membrane proteins. Collectively, these results reveal how the σ^M^ regulon has evolved to up-regulate membrane-localized complexes involved in cell wall synthesis, and to simultaneously counter the resulting stresses imposed by regulon induction.

**Author Summary:** Bacteria frequently produce antibiotics that inhibit the growth of competitors, and many naturally occurring antibiotics target cell wall synthesis. In *Bacillus subtilis*, the alternative σ factor σ^M^ is induced by cell wall antibiotics, and upregulates genes for peptidoglycan and cell envelope synthesis. However, dysregulation of the σ^M^ regulon, resulting from loss of the YhdL anti-σ^M^ protein, is lethal. We here identify charge variants of the SpoIIIJ(YidC) membrane protein insertase that suppress the lethal effects of high σ^M^ activity. Further analyses reveal that induction of the σ^M^ regulon leads to high level expression of membrane proteins that trigger envelope stress, and this stress is countered by specific genes in the σ^M^ regulon.

## Introduction

The ability of cells to adapt to changing conditions relies in large part on the expression of specific stress responses controlled by transcription regulators. The extracytoplasmic function (ECF) subfamily of σ factors are frequently involved in bacterial responses to stresses affecting the cell envelope [1, 2]. In *Bacillus subtilis*, there are seven ECF σ factors with the σ^M^ regulon playing a particularly important role in modulating pathways involved in peptidoglycan synthesis, cell envelope stress responses, and intrinsic resistance to antibiotics [3]. Cells lacking σ^M^ (*sigM* null mutants) grow normally in unstressed conditions, but have greatly increased sensitivity to high salt and to cell wall active antibiotics, including ß-lactams and moenomycin [4–6].

Like many other ECF σ factors, the activity of σ^M^ is regulated by two membrane-localized anti-σ factors encoded as part of the *sigM* operon, *sigM-yhdL-yhdK* [7, 8]. The major anti-σ factor is YhdL, a transmembrane protein that directly binds σ^M^ [9]. However, full YhdL activity requires a second transmembrane protein YhdK. Although a lack of σ^M^ is well tolerated under unstressed conditions, the lack of the anti-σ^M^ factors leads to a runaway activation of the autoregulated *sigM* operon, and overexpression of the σ^M^ regulon [10]. A null mutation of *yhdK* leads to an ∼100-fold elevation of σ^M^ activity, morphological abnormalities, and slow growth [10]. A null mutation in *yhdL* is lethal, but suppressors arise readily that have inactivated *sigM* and grow normally in unstressed conditions [4, 10]. These findings imply that high level expression of σ^M^ is toxic to cells, but the basis for this toxicity has not been defined.

Previously, we explored the basis for σ^M^ toxicity by selecting for suppression of *yhdL* lethality in a *sigM* merodiploid strain to reduce the frequency of suppressors that had inactivated σ^M^. These studies led to the recovery of mutations in *rpoB* and *rpoC*, encoding the ß and ß’ subunits of RNA polymerase, that led to a reduction of σ^M^ activity sufficient to restore viability [10]. These mutations, which acted selectively on σ^M^, affected a region of core RNA polymerase involved in σ factor binding. In the course of these studies we also demonstrated that the toxicity from high σ^M^ could be alleviated by mutation of the autoregulatory promoter for the *sigM* operon, or by overexpression of the housekeeping σ factor, σ^A^ [10]. These results suggest that the lack of a functional anti-σ^M^ factor (*yhdL* null mutant) leads to runaway activation of the *sigM* operon and a high level of σ^M^ activity that is incompatible with growth. However, it is unclear whether σ^M^ toxicity results from a decrease in activity of the essential housekeeping σ^A^, overexpression of one or more σ^M^-regulated genes, or both.

To further explore the impact of overexpression of specific σ^M^-regulated genes on cell physiology we generated a library of strains in which specific σ^M^-dependent promoters (P_M_) are inactivated by point mutations. This approach, which removes σ^M^-dependent activation while leaving other promoters and regulatory inputs intact, is important since many σ^M^-regulated genes have multiple promoters and encode essential genes, including several involved in peptidoglycan synthesis and cell division [3]. In the course of developing this library we received a previously described strain with a mutation inactivating the P_M_ of *rodA*, encoding a SEDS family transglycosylase important for peptidoglycan synthesis [6]. We unexpectedly discovered that in this strain *yhdL* could be inactivated, and the same was true for the parent strain (*B. subtilis* PY79). These serendipitous observations led us to hypothesize that *B. subtilis* PY79 differs from other *B. subtilis* 168 strains in its ability to tolerate high level expression of the σ^M^ regulon.

Here, we report that the ability of *B. subtilis* PY79 to tolerate σ^M^ regulon overexpression results from a single amino acid substitution in the *spoIIIJ* gene, which encodes the major YidC membrane insertase in *B. subtilis* [11, 12]. This finding motivated a detailed structure-function analysis of YidC, which led to the discovery of mutations that increase the positive charge within the hydrophilic, substrate-binding channel of YidC from +1 (wild-type) to +2 or +3 (in specific combinations) increase tolerance to overexpression of σ^M^-regulated membrane proteins. Moreover, high level activity of σ^M^ leads to induction of genes associated with secretion stress, oxidative stress, and DNA damage responses, and the σ^M^ regulon itself includes functions that help compensate for stresses associated with membrane protein overexpression.

## Results

### A single amino acid change in SpoIIIJ necessary and sufficient for tolerance of high **σ**^M^

The anti-σ factors YhdL and YhdK regulate σ^M^, and the absence of *yhdL* is lethal in *B. subtilis* strain 168 due to toxic levels of σ^M^ [4, 10]. Since σ^M^ controls a large regulon, including many essential genes involved in cell wall synthesis [1, 3], we sought to construct a library of strains in which specific σ^M^-dependent promoters are inactivated by point mutations. One such promoter precedes *rodA*, which encodes a peptidoglycan transglycosylase [6]. We thus acquired a ΔP_M_-*rodA* strain (BAM1077[6]), and tested whether *yhdL* is essential in that strain background, with its parent wild type strain PY79 as a control. Surprisingly, we found that *yhdL* is not essential in either PY79 strain. A *yhdL* mutant in PY79 exhibits reduced colony size compared to WT and high σ^M^ activity as indicated by the blue color on LB plates containing X-Gal (Fig. 1a). This mutant is relatively stable, with the occasional appearance of suppressors that have a large white colony morphology (likely containing mutations in *sigM*). In contrast, introduction of a *yhdL* null mutation into other *B. subtilis* 168 strains results in tiny, pinpoint colonies that cannot be re-streaked, consistent with prior work [4, 10].

**Fig. 1.**
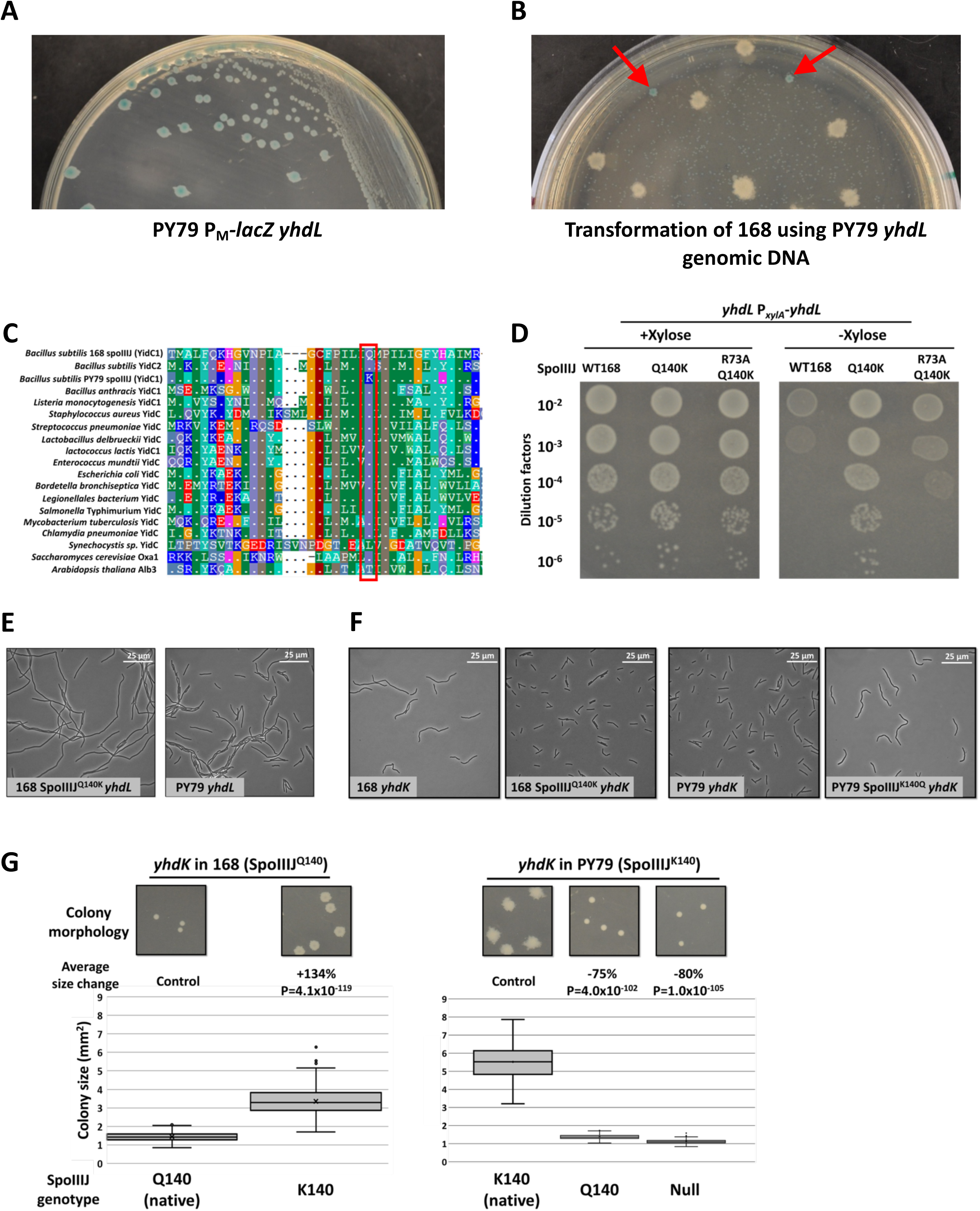
Identification of a SpoIIIJ variant that is necessary and sufficient to abolish the essentiality of the anti-σ^M^ factor YhdL. (A) Growth of PY79 P_M_-*lacZ yhdL* strain on LB plate with 60 μg/ml X-Gal. (B) Transformation plate using 168 P_M_-*lacZ* as recipient and PY79 P_M_-*lacZ yhdL*::*kan* as DNA donor, with LB medium supplemented with kanamycin and X-Gal. Red arrows are pointed at intermediate sized blue colonies that were further analyzed by whole genome re-sequencing. (C) Alignment of YidC homologs from bacterial and eukaryotic species. Region of the Q140 of *B. subtilis* SpoIIIJ is shown. (D) Spot dilution of *yhdL* P*_xylA_*-*yhdL* depletion strain with different SpoIIIJ alleles. WT168 has SpoIIIJ^Q140^ allele. (E) Representative phase contrast microscopy image of *yhdL* mutants in different strain backgrounds. PY79 WT has SpoIIIJ^K140^ allele. (F) Representative phase contrast microscopy photo of *yhdK* mutants in different strain backgrounds. (G) Colony morphology and size of *yhdK* mutants in different backgrounds. Colony size data were plotted using Box and Whisker chart and the bottom and top of the box are the first and third quartiles, the band inside the box is the second quartile (the median), and the X inside the box is the mean. Whiskers are one standard deviation above or below the mean. Outliers are shown as single dots. Average colony size change was calculated using formulation “change = (sample - control) / control x 100%”. P value was calculated using student’s t test, two tails, assuming unequal variances. Sample number n is at least 100 for each sample.

To identify the genetic differences in PY79 that confer tolerance of high σ^M^, we compared the genome between *B. subtilis* strain 168 and PY79. There are over a hundred single nucleotide polymorphisms (SNPs) between the 168 reference sequence and PY79, as well as four large deletions in the genome of PY79 (including the SPβ prophage), causing a reduction of 180 kb for the PY79 genome compared with 168 [13, 14]. To identify differences that might correlate with tolerance of high σ^M^ activity, we compared the sequences of genes encoding RNA polymerase subunits, and genes in the σ^M^ and Spx regulons (Spx is a transcription regulator that is regulated by σ^M^). However, no differences were noted in these 103 genes between the PY79 and 168 reference genomes (Table S1).

Next, we used an unbiased, forward genetics approach to identity the mutation in PY79 that suppresses the lethality of a *yhdL* null mutation. We used genomic DNA from a PY79 *yhdL*::*kan* strain to transform 168 to kanamycin resistance, reasoning that the only viable transformants will likely also acquire the suppressing mutation. Because each competent cell of *B. subtilis* contains about 50 binding sites for DNA uptake, a competent cell can import multiple fragments of DNA during transformation in a process known as congression [15]. When a 168 strain containing a P_M_-*lacZ* reporter was transformed with chromosomal DNA from the viable PY79 *yhdL*::*kan* strain and selected on an LB plate supplemented with kanamycin and X-gal, we recovered numerous tiny blue colonies that did not grow when re-streaked onto fresh plates (consistent with the essentiality of YhdL in the 168 background), a few large white colonies (likely *sigM* mutants), and intermediate sized blue colonies (Fig. 1B). The intermediate blue colonies grew to a similar size as a *yhdL* null mutant in PY79, consistent with acquisition of both the *yhdL*::*kan* allele and a second locus that suppresses the toxicity of high σ^M^. Whole genome sequencing was performed on these transformants and the reads were mapped to the reference genome of 168. Out of 15 sequenced 168 transformants, 14 contained the same SNP imported from PY79 that generates a missense mutation in *spoIIIJ* (encoding SpoIIIJ^Q140K^) (Fig. S1A, Table S2). We therefore hypothesized that the *spoIIIJ*^Q140K^ allele was the suppressor needed for cells to tolerate the *yhdL* mutation.

SpoIIIJ belongs to the YidC membrane protein insertase family and is responsible for inserting membrane proteins into the lipid bilayer, independently or in association with the Sec secretion system [16–18]. *E. coli* encodes one essential homolog of YidC, while some bacteria such as *B. subtilis* encode two homologs, SpoIIIJ(YidC) and YidC2 [18, 19]. The gene encoding YidC was named *spoIIIJ* because mutations at this locus lead to a block at stage III of sporulation [19, 20]. However, *spoIIIJ* is constitutively expressed and functional in vegetative cells. The expression of the paralog YidC2 is regulated by an upstream gene *mifM*, which monitors the total membrane protein insertase activity and only allows expression of YidC2 when MifM is not efficiently inserted into the membrane [21]. Both SpoIIIJ and YidC2 can fulfill the essential function of YidC insertase, with SpoIIIJ essential for sporulation [22] and YidC2 important for the development of competence (Fig. S2A) [23]. Interestingly, an alignment of YidC homologs revealed that the Gln140 residue is highly conserved among bacteria, and only *B. subtilis* PY79 SpoIIIJ contains Lys at this position (Fig. 1C).

To test if this SpoIIIJ Gln to Lys variant (SpoIIIJ^Q140K^) is necessary and sufficient for tolerance of high σ^M^, we introduced the *spoIIIJ*^Q140K^ mutation at the native locus of strain 168 using CRISPR, and found that *yhdL* was no longer essential (Fig. 1D, Fig. S1B). Conversely, changing the Lys140 into Gln in PY79 abolished the ability of PY79 to tolerate loss of *yhdL* (Fig. S1B), suggesting that the SpoIIIJ^Q140K^ is necessary and sufficient for tolerance of a *yhdL* deletion mutation. To test if the *spoIIIJ*^K140^ allele is dominant over the *spoIIIJ*^Q140^ allele, we constructed merodiploid strains expressing both alleles of SpoIIIJ (using a vector with xylose-inducible promoter P*_xylA_* that integrates into *ganA* and when induced produces about 70% amount of the native protein level (Fig. 2B)). Strains with either a P*_xylA_*-*spoIIIJ*^K140^ in the 168 strain, or P*_xylA_*-*spoIIIJ*^Q140^ in PY79 strain background could still tolerate the loss of *yhdL* (Fig. S1B). This dominance suggests that SpoIIIJ^Q140K^ leads to a gain of function that enables cells to tolerate high σ^M^ activity. Phase contrast microscopy revealed that a 168 SpoIIIJ^Q140K^ *yhdL* mutant had a similar but slightly more elongated cell morphology compared with a PY79 *yhdL* mutant (Fig. 1E), confirming the major role of SpoIIIJ^Q140K^ in tolerance of a *yhdL* null mutation.

**Fig. 2.**
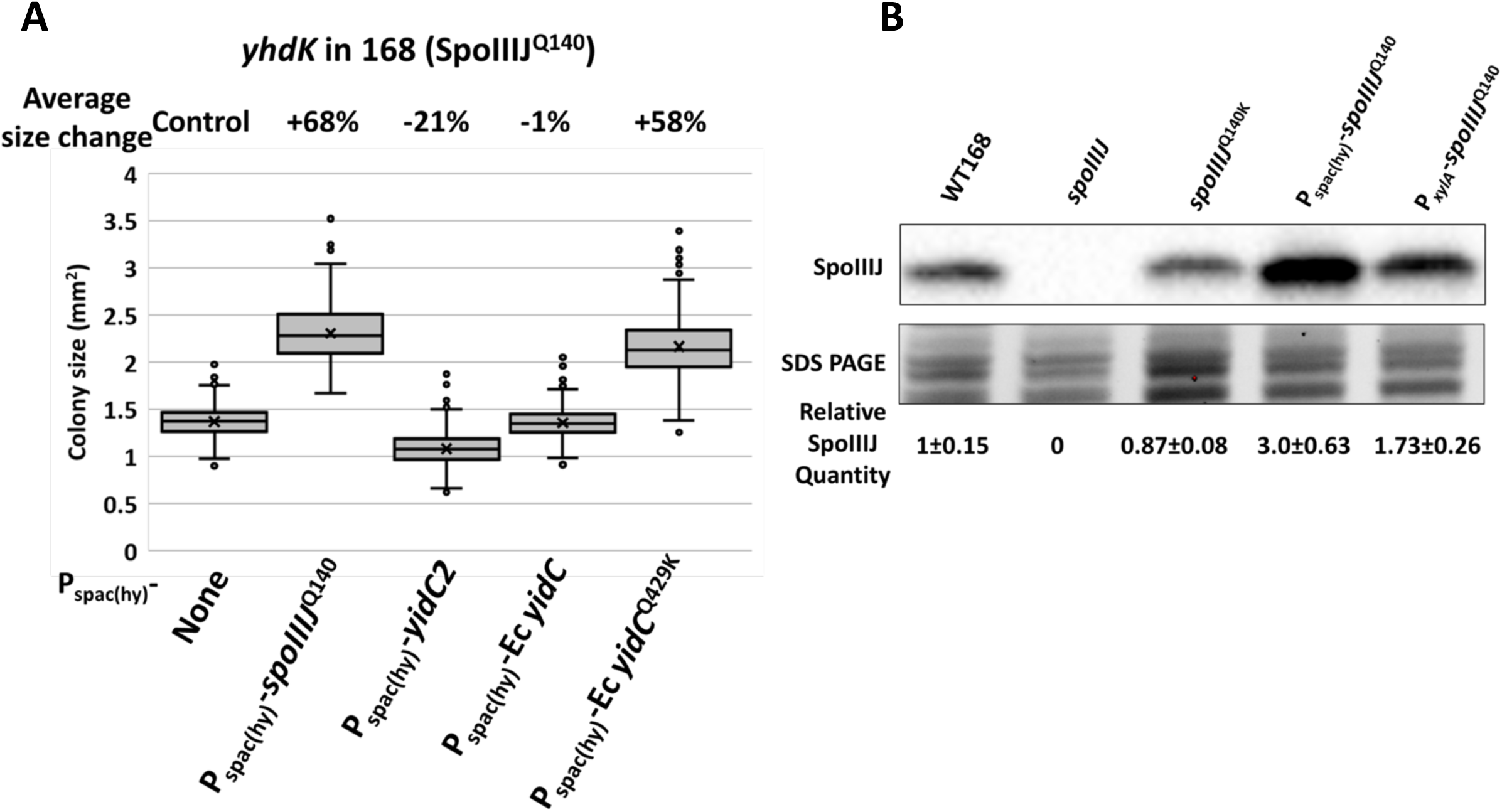
Overexpression of SpoIIIJ increases tolerance of high σ^M^ activity. (A) Colony size of *yhdK* mutants in 168 background with overexpression of YidC homologs using an IPTG inducible P_spac(hy)_ promoter. Cells were grown on LB plate supplemented with 1 mM IPTG for 24 hours at 37 °C. Box and Whisker chart was plotted as in Fig. 1G. (B) Representative Western blot of SpoIIIJ protein in different strain backgrounds. P_spac(hy)_ and P*_xylA_* induced SpoIIIJ strains contain the native SpoIIIJ and were induced with 1 mM IPTG or 1% (w/v) xylose, respectively. Part of the SDS PAGE gel was shown as a loading control. Relative SpoIIIJ quantity was calculated using band intensity values from three biological replicates and normalized using total lane intensity from the SDS PAGE gel. Data is presented as mean ± SEM.

As a general strategy to monitor cell fitness, we compared the impact of *spoIIIJ* alleles on morphology and colony size in the PY79 and 168 backgrounds mutant for *yhdK*. YhdK functions together with YhdL as an anti-σ complex, but unlike *yhdL* a *yhdK* null mutant is tolerated in 168 strains, although it does lead to an ∼100-fold increase in σ^M^ regulon expression and severe growth defects [10]. Five independent trials with a 168 *yhdK* mutant, known to generate small colonies, revealed up to a ∼10% change in average colony area. This variation likely results from small differences in the plating conditions. Although these small differences were in some cases judged to be statistically significant (based on P value in a t test, two tails, assuming unequal variances) (Fig. S1C), this reflects the high sample number in each measurement (100-1000 colonies per measurement). Considering this level of variation between genetically identical strains, we only regard as significant those changes of >10% in colony size. Using this assay, we found that the small colony size, as well as the filamentous cell morphology of the 168 *yhdK* mutant, can be largely rescued by the *spoIIIJ*^Q140K^ allele (Fig. 1F, G). Conversely, a *spoIIIJ*^K140Q^ mutation in PY79 *yhdK* converted the large colonies of the parent strain into the small round morphology of the 168 *yhdK* strain, and the cells exhibited increased filamentation (Fig. 1F, G). Deletion of *spoIIIJ* in a PY79 *yhdK* mutant mimicked a SpoIIIJ^K140Q^ mutation (Fig. 1G), likely because the cells now rely on the other YidC paralog, YidC2, which contains a glutamine residue in the equivalent position (Fig. 1C) [19]. Overall, our results show that SpoIIIJ^Q140K^ mutation is necessary and sufficient for *B. subtilis* to tolerate high σ^M^ activity caused by the absence of the anti-σ factor YhdL or its partner protein YhdK.

### Overexpression of SpoIIIJ increases tolerance of high σ^M^ activity

We hypothesized that the SpoIIIJ^Q140K^ protein may simply be more active or abundant in cells than the native protein. To test if increasing insertase activity is sufficient to alleviate toxicity associated with high σ^M^, we overexpressed the wild-type 168 SpoIIIJ^Q140^ protein using a strong IPTG inducible promoter, P_spac(hy)_ [24]. Induction of the P_spac(hy)_-*spoIIIJ*^Q140^ allele led to a three-fold increase in the amount of SpoIIIJ protein compared with WT (Fig. 2B), and increased the colony size of a *yhdK* mutant by 68% (Fig. 2A). This increase is less than the effect of the *spoIIIJ*^Q140K^ allele at the native locus (which increased *yhdK* colony size by 134%), and consistently it only marginally increased the growth of a *yhdL* depletion strain under depletion condition (Fig. S2C). The increase in the fitness of the *yhdK* mutant supports the hypothesis that higher insertase activity is beneficial for cells with elevated σ^M^ activity, but overexpression alone does not phenocopy the effect of the altered function allele.

We next tested whether overexpression of YidC2, the other YidC homolog in *Bacillus*, could benefit cells with high σ^M^ expression. Interestingly, when *yidC2* was overexpressed from the P_spac(hy)_ promoter, the growth defect of either the *yhdK* mutant or the *yhdL* depletion strain was exacerbated (Fig. 2A, S2C). Furthermore, when the equivalent glutamine residue of SpoIIIJ^Q140^ was mutated to lysine, the YidC2^Q148K^ mutant protein was toxic when overexpressed in a *yhdK* mutant (Fig. S2B). Thus, YidC2 is unable to compensate for SpoIIIJ in alleviating stress associated with high σ^M^ activity, even when the corresponding Gln to Lys substitution is present. Similarly, overexpression of *E. coli* YidC did not provide any benefit to a *yhdK* or *yhdL* mutant (Fig. 2A). However, when the equivalent Gln to Lys substitution was present, overexpression of the *E. coli* YidC^Q429K^ mutant was modestly beneficial, and colony size of the *yhdK* mutant increased by 58% (Fig. 2A). These results suggest that different YidC homologs vary in their substrate preferences, and the Gln to Lys mutation may enhance the ability of the *B. subtilis* SpoIIIJ and *E. coli* YidC insertases to facilitate membrane insertion of at least some σ^M^-dependent proteins.

### SpoIIIJ^Q140K^ increases the positive charge inside the substrate binding groove

The structure of YidC2 from *Bacillus halodurans* revealed a positively charged hydrophilic groove formed by five transmembrane segments [16]. A positively charged residue in the substrate-binding groove is essential for the function of the insertase, as a R73A substitution in *B. subtilis* SpoIIIJ completely abolished the essential function of SpoIIIJ *in vivo,* while an R73K substitution retained function [16]. Although the positive charge is essential, the R73 residue is not, as the positive charge can be provided by mutation of any of six other residues inside the hydrophilic groove to Arg. These six positions include Ile72 and Ile76 in transmembrane region 1 (TM1), Gln140 and Leu144 in TM2, and Trp228 and Gly231 in TM5 [25] (Fig. 3A, S3B). Since both Arg and Lys are positively charged, we hypothesized that the key feature of the PY79 SpoIIIJ^K140^ protein is the +2 charge inside its substrate binding chamber. To test this hypothesis, we engineered a SpoIIIJ^R73AQ140K^ double substitution protein with a net charge of +1, where R73 is functionally replaced by K140. Expression of this protein is sufficient to support viability, as judged in a strain with depletion of *yidC2* (Fig. 3B), but is not sufficient to allow depletion of *yhdL* (Fig. 1D). This supports the idea that the key effect of the Q140K substitution is to increase the positive charge in the substrate binding groove.

**Fig. 3.**
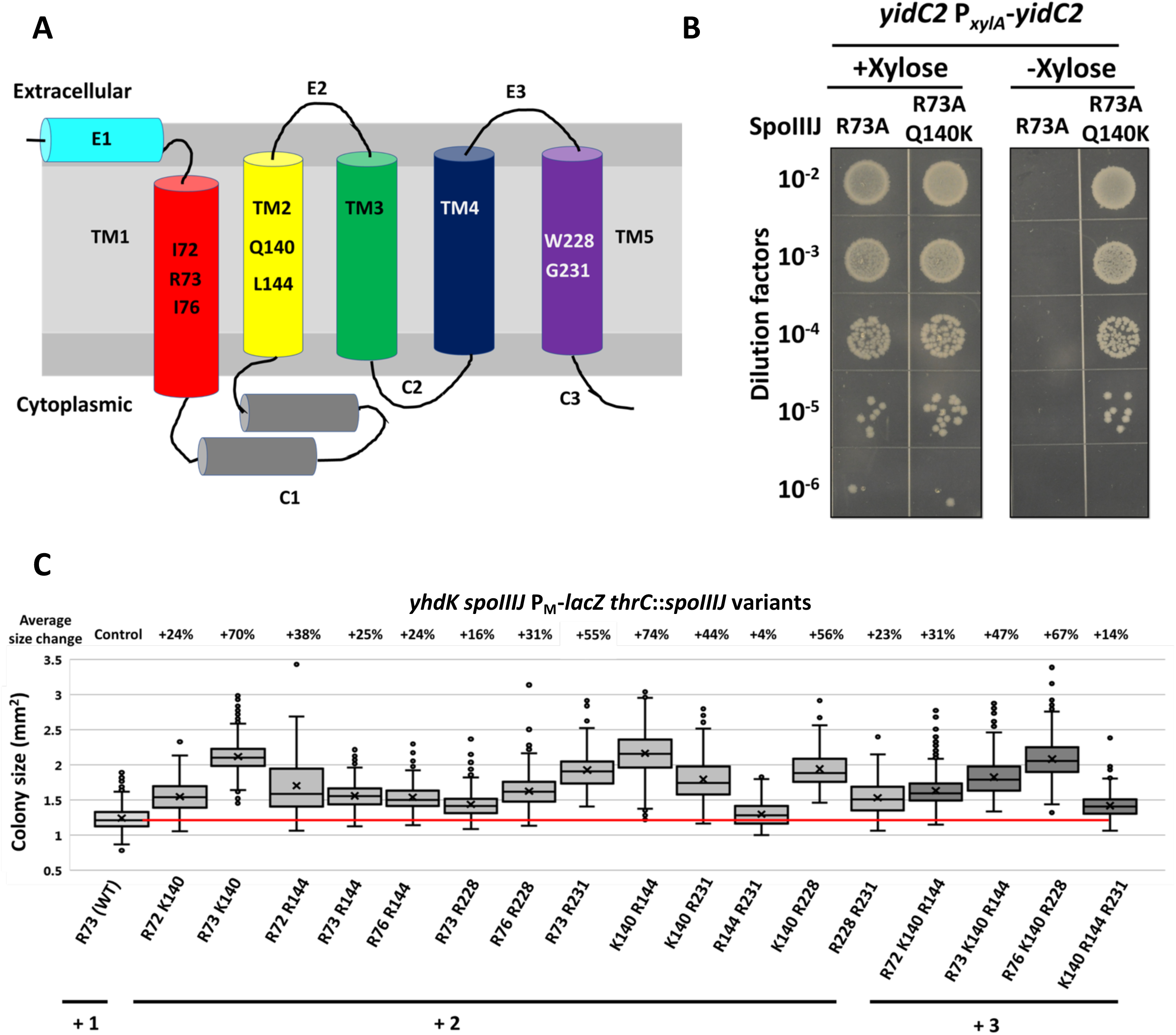
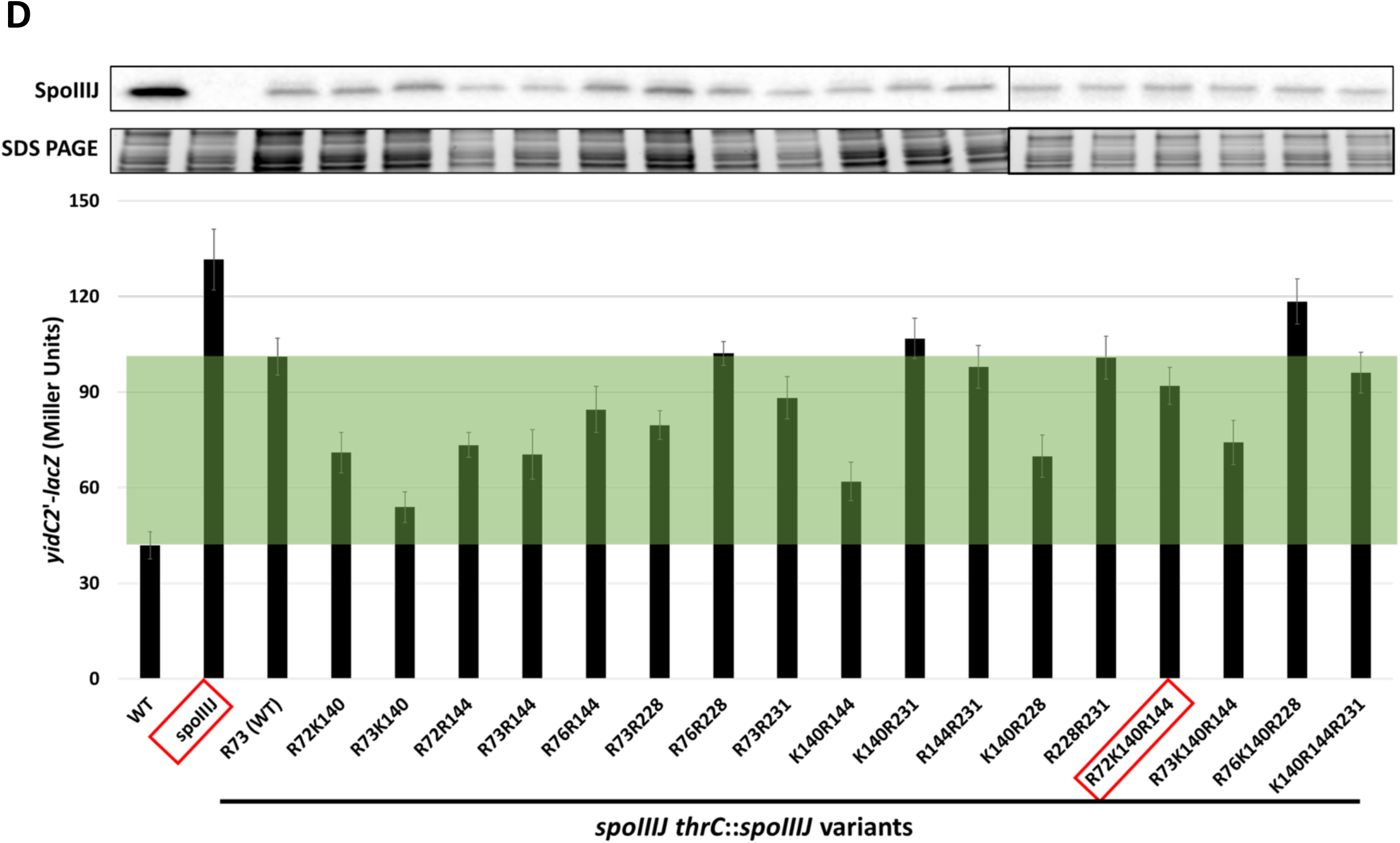
Effects of different charge variants inside the SpoIIIJ substrate binding groove. (A) Schematic drawing showing the extracellular, transmembrane and cytoplasmic domains of SpoIIIJ. The native sequence in strain 168 of the seven residues for the positive charge library were shown. (B) Spot dilution of *yidC2* P*_xylA_*-*yidC2* depletion strain with different SpoIIIJ variants in strain 168. (C) Colony size of *yhdK spoIIIJ* mutants with different *spoIIIJ* alleles integrated in *thrC* site. Residues that provide positive charge(s), as well as total positive charge(s) provided by these seven variable positions are labelled. The red line shows the median of colony size with a WT SpoIIIJ from strain 168. Box and Whisker chart was plotted as in Fig. 1G. (D) Top, representative Western blots of SpoIIIJ protein in different strain backgrounds. Part of the SDS PAGE gels were shown as loading control. Bottom, activity of *yidC2*’-*lacZ* translational fusion in the same strain backgrounds of the Western blots. “WT” has a native copy of *spoIIIJ* while “*spoIIIJ*” is a markerless deletion mutant. All other strains have the native *spoIIIJ* deleted and contain variants of *spoIIIJ* at *thrC* site. Among the seven variable positions, residues that provide positive charge(s) are labelled. SpoIIIJ variants inside the red boxes cannot support growth of a *yidC2* P*_xylA_*-*yidC2* depletion strain in the absence of inducer xylose. Green shading highlights the difference between the activity of SpoIIIJ at the native and *thrC* loci. Variants with *yidC2*’-*lacZ* activity inside the green shading exhibit increased MifM insertion activity compared with WT.

### Increased charge inside the substrate binding groove is key for tolerance of high σ^M^ activity

Since it is the increased positive charge from +1 to +2, rather than the Q140K substitution *per se*, that rescues cells from high σ^M^ toxicity, we next set out to test if other combinations of positively charged residues can also rescue cells from high σ^M^ activity. To this end, we generated a library of SpoIIIJ variants with five of the six positions mentioned above substituted (or not) with Arg, as shown previously to support function in the absence of Arg73 [25], and Gln140 substituted (or not) with Lys, as seen in PY79. In addition, we mutated Arg73 to Ala (or not) (Fig. 3A). This leads to 2^7^=128 possible charge combinations, ranging from 0 to a maximum of +7, with most containing a nominal positive charge of +2 to +5 in the substrate binding groove (Fig. S3A, S3B). Note that for the sake of simplicity, we assume that the K140 residue is positively charged, since the epsilon-amino group of free Lys has a pKa of ∼10.5, but this will likely vary depending on the local charge environment.

To identify SpoIIIJ variants that can function to support growth of a *yhdL* depletion strain, we transformed a *spoIIIJ* null *yhdL* depletion strain and selected for transformants that grew in the absence of *yhdL* induction. After sequencing and validation, we identified 103 *spoIIIJ* mutants that supported growth in the absence of *yhdL* induction (Table S3). Among them, 88 mutants contained a nominal double positive charge in 13 unique combinations (Fig. S3A). Five of these 13 combinations include K140, including the combination present in the PY79 SpoIIIJ protein: R73 K140. The remaining 15 mutants contained a nominal triple positive charge, with four unique combinations. Interestingly, all four of these variants contained K140, and in each case K140 was present together with a pair Arg residues that also supported growth with Q140. It is possible that the presence of nearby Arg residues lowers the pKa of K140, and the protonation state of this residue may also vary depending on the nature of the bound substrate. Among the 17 unique combinations of double and triple positive charges, none has more than one positive charge in TM1, whereas TM2 and TM5 can each harbor double positive charges (Fig. S3A, Table S3). We conclude that all 17 functional SpoIIIJ variants have an effective charge of between +2 and +3 in the substrate binding channel. This strongly suggests that a modest increase of charge inside the substrate binding chamber facilitates the insertion of σ^M^-regulated proteins overproduced under YhdL depletion conditions, whereas a further increase may be detrimental to the activity or the stability of the insertase.

Each of these 17 SpoIIIJ variants can alleviate the stress imposed by high σ^M^ activity and support growth of the YhdL depletion strain (Fig. S3C), and increase the colony size of the *yhdK* mutant by up to 74% (Fig. 3C). We also found 16 of these 17 variants can support growth of a YidC2 depletion strain (Fig. S3D). Only the SpoIIIJ R72 K140 R144 variant was unable to support cell growth under these conditions, suggesting that it is compromised in the ability to insert proteins essential for cell growth, despite its ability to modestly increase colony size of the *yhdK* mutant (31%).

SpoIIIJ also inserts MifM into the membrane, which serves as a sensor of SpoIIIJ function to regulate expression of *yidC2* [21, 26]. We used a *yidC2’*-*lacZ* translational fusion reporter to measure the ability of each SpoIIIJ variant to insert MifM. A WT 168 strain exhibited very low level of *yidC2*’-*lacZ* activity in the presence of native *spoIIIJ* expression (∼42 Miller Units (MU), Fig. 3D), and deletion of *spoIIIJ* increased the reporter activity by about 3-fold (∼132 MU, Fig. 3D). Complementation of a *spoIIIJ* null mutant with the 168 version of *spoIIIJ* (single positive charge at R73) at the *thrC* locus reduced the reporter activity to ∼101 MU (Fig. 3D). The lack of complete complementation may be caused by the location of the gene, as the native locus is close to the origin of the chromosome and thus has higher copy number than *thrC* in fast growing cells. Indeed, under our growth condition (late exponential phase in LB at 37°C), less SpoIIIJ protein was detected in the *thrC*::*spoIIIJ* complementation strain than the WT (Fig. 3D).

Using this strain background, we found that most of the SpoIIIJ variants with double positive charge appear to have higher MifM insertion ability as they exhibited lower *yidC2’*-*lacZ* activity than the WT protein. Interestingly, the three mutants with double positive charge that have the largest colony size in a *yhdK* mutant background (R73 K140, K140 R144, and K140 R228), also showed the lowest *yidC2’*-*lacZ* activity, consistent with the hypothesis that they have higher insertase activity for both MifM and for proteins that contribute to toxicity in strains with high σ^M^-activity. Conversely, one mutant with a nominal triple positive charge (R76 K140 R228) exhibited a strong increase in colony size in the *yhdK* mutant background (Fig. 3C), but appeared to have low MifM insertion activity (high *yidC2’-lacZ* expression; Fig. 3D). Overall these results suggest that increasing the positive charge inside the substrate binding groove affects the efficiency of inserting membrane proteins, with several variants that appear to enhance the insertion efficiency for both MifM and σ^M^-regulated proteins, while also retaining the ability to insert essential membrane proteins.

### High σ^M^ activity causes a cascade of stresses that can be partially compensated by its regulon

SpoIIIJ functions as a membrane insertase, by either independently inserting some single pass membrane proteins or by functioning as part of the Sec translocon to facilitate the folding of the translocated proteins [18, 27]. The finding that SpoIIIJ^Q140K^ can suppress growth defects caused by high σ^M^ leads us to reason that the toxicity of high σ^M^ is caused by overexpression of membrane proteins that overwhelm the secretion system, and that SpoIIIJ^Q140K^ suppresses this stress by functioning as a more efficient insertase/foldase or chaperon. In *B. subtilis*, membrane secretion stress is sensed by the CssRS (<u>c</u>ontrol of <u>s</u>ecretion <u>s</u>tress <u>r</u>egulator/<u>s</u>ensor) two component system. In the presence of membrane secretion stress, caused either by overproduction of secreted proteins or high temperature, CssR upregulates expression of membrane proteases HtrA (<u>h</u>igh temperature <u>r</u>equirement A) and HtrB to facilitate the re-folding or degradation of misfolded proteins [28, 29].

To test whether high σ^M^ causes secretion stress, we constructed a P_*htrA*_-*lux* reporter to monitor induction of the CssR-regulon. Indeed, P*_htrA_* activity was increased about ten-fold in a *yhdK* mutant, and this induction can be reduced by the SpoIIIJ^Q140K^ allele by about 70% (Fig. 4A). To evaluate the role of the CssRS system in alleviating the membrane protein overproduction stress under high σ^M^ condition, we constructed mutants missing components of the CssR regulon and used *yhdK* mutant colony size as a measure of fitness. Surprisingly, we found that neither CssR-regulated membrane protease (HtrA and HtrB) is important for fitness of the *yhdK* mutant as judged by the effect of their deletion on colony size. Deletion of the response regulator CssR abolished the induction of the regulon [29], and has no effect on the fitness of *yhdK* mutant. In contrast to a previous report [29], the absence of CssS increased transcription of P*_htrA_*-*lux* reporter by ∼80-fold. This suggests that CssS may act both as a sensor kinase to activate CssR and the regulon under induction conditions, and also as a phosphatase to deactivate CssR in the absence of induction signals, as also noted for sensor kinases in other two component systems [30].

**Fig. 4.**
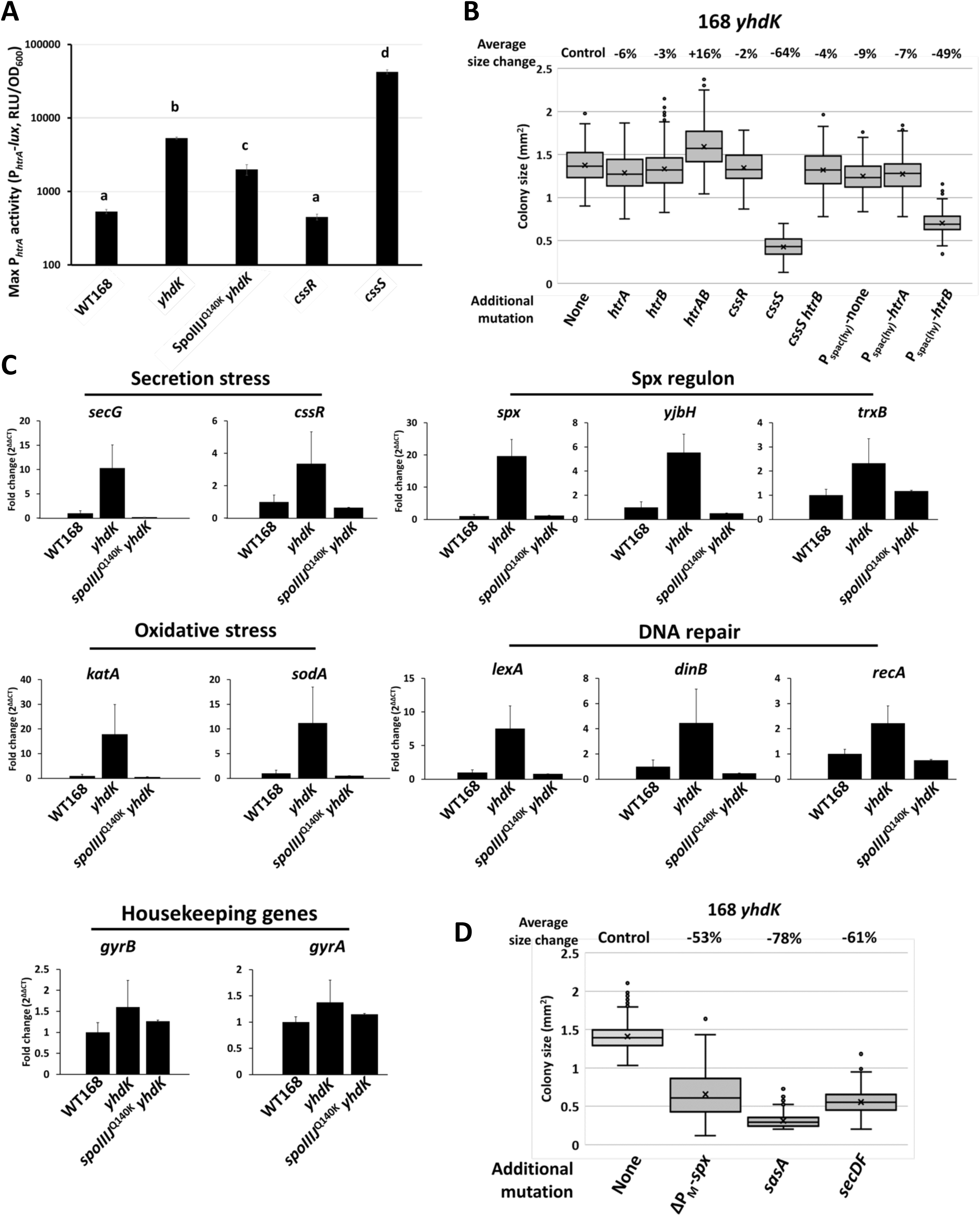
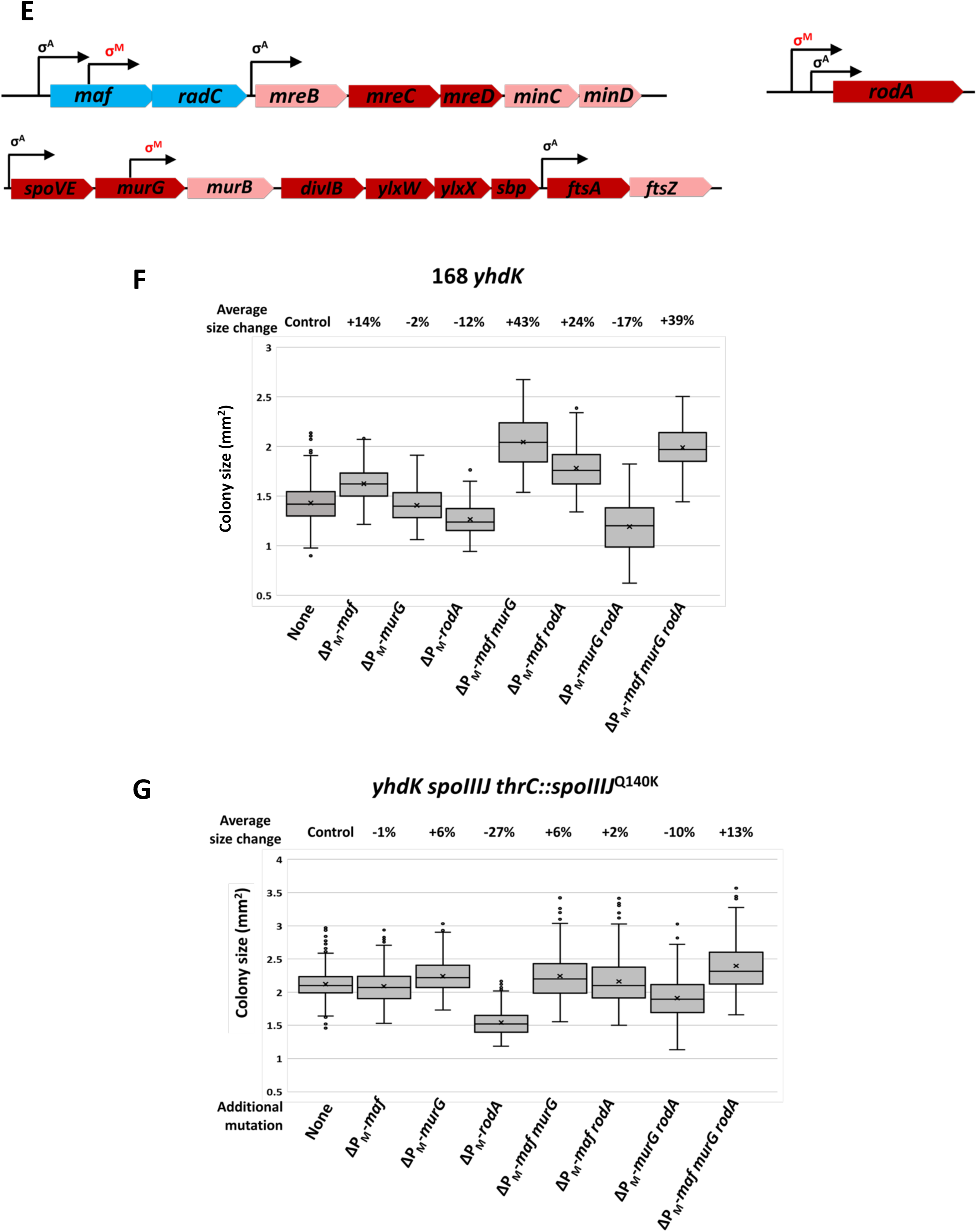
High σ^M^ activity causes a cascade of stresses that can be partially compensated by its own regulon. (A) Maximum promoter activity of *htrA* measured using a P*_htrA_*-*lux* transcriptional fusion reporter in different strain backgrounds. Data is presented as mean ± SEM, and statistically significant different samples (Student’s t test, two-tailed P<0.05) are labelled with different letters. Sample number n is at least 4. (B) Colony size of *yhdK* mutants in strain 168 with different additional mutations. Box and Whisker chart was plotted as in Fig. 1G. (C) Fold change of transcription levels of representative genes for stress response or housekeeping genes. Data is presented as mean ± SEM. Sample number n is at least 4. (D) Colony size of *yhdK* mutants in strain 168 with different additional mutations. ΔP_M_-*spx* strain has the native *spx* deleted and an ectopic copy without the P_M_ promoter [42]. (E) Schematic drawing of *maf, murG* operon and *rodA* gene. σ^A^ and σ^M^ controlled promoters are shown with arrows. Cytoplasmic proteins are labelled in blue boxes, peripheral membrane proteins in pink boxes, and membrane integral proteins in red boxes. (F, G) Colony size of *yhdK* mutants with different P_M_ promoter(s) mutated in (F) WT strain 168 or (G) strain 168 with the native *spoIIIJ* deleted and a SpoIIIJ^Q140K^ allele integrated at the *thrC* site. Box and Whisker chart was plotted as in Fig. 1G.

Deletion of *cssS* reduced *yhdK* mutant colony size by over 60% (Fig. 4B), and the reduction of colony size is reversed in a *cssS htrB* double mutant, suggesting that high HtrB activity is detrimental for the *yhdK* mutant. Consistently, when *htrB* but not *htrA* was overexpressed using the P_spac(hy)_ promoter, the colony size of *yhdK* mutant was reduced by about 50% (Fig. 4B). Deletion of these genes in the WT 168 background did not have much effect on the colony size (Fig. S4A). These results suggest that the *yhdK* mutant is sensitive to the overproduction of membrane proteases. Single deletion of other proteases, including SipT, SipS, HtpX, PrsW and GlpG in a *yhdK* null mutant was also not detrimental, and in some cases led to a small beneficial effect in colony size (Fig S4B), again indicating that membrane protease activity is not beneficial for the *yhdK* mutant. We speculate that under high σ^M^ conditions, the overproduced proteins may cause a backlog of membrane proteins that require a longer time to correctly insert and fold in the membrane. Some of these proteins may be essential for cell growth and are vulnerable to degradation by membrane proteases.

Misfolding of membrane proteins, as well as jamming the Sec translocon by hybrid proteins, leads to the generation of reactive oxygen species (ROS) and ultimately DNA damage [31, 32]. To test whether high σ^M^ also triggers a similar cascade of stresses, we performed quantitative RT-PCR to measure induction of representative stress genes. Indeed, the high σ^M^ activity in the *yhdK* null mutant was correlated with strong induction of genes involved in secretion stress (*secG*, *cssR*), oxidative stress (*katA*, *sodA*), and DNA repair (*lexA*, *dinB* and *recA*) (Fig 4C). The induction of these stress response genes was largely suppressed in the presence of the s*poIIIJ*^Q140K^ allele, suggesting that their induction is a downstream effect resulting from inefficient insertion of membrane proteins (Fig 4C).

Interestingly, while many genes in the σ^M^ regulon are involved in the synthesis and maintenance of the cell wall, there are also genes involved in regulation of redox balance (*spx* and the Spx regulon), DNA repair and recombination (for example, *radA*, *radC* and *recU*), and ppGpp synthesis and stringent response (*sasA*) [3]. There is also a candidate σ^M^ promoter associated with the *secDF* operon [3]. This suggests that a subset of genes in the σ^M^ regulon are involved in compensating for stresses associated with upregulation of σ^M^. Indeed, deletion of *secDF*, *sasA* or the P_M_ of *spx* dramatically reduced the colony size of the *yhdK* mutant (Fig. 4D), while there was very little effect noted in the WT background (Fig. S4C). Thus, these genes seem to play an important role in cell fitness specifically under conditions of high σ^M^ expression. Deleting single genes inside the Spx regulon in the *yhdK* mutant did not lead to noticeable reduction of colony size (Fig. S4D), suggesting functional redundancy within the Spx regulon.

Among the 69 genes currently assigned to the σ^M^ regulon according to SubtiWiki [33], 38 code for membrane-associated or secreted proteins (Table S4). To demonstrate the burden these membrane proteins may cause when overexpressed under high σ^M^ condition, we mutated the σ^M^-dependent promoters (P_M_) for genes important for the elongasome and the divisome and tested the consequence on fitness measured by colony size of the *yhdK* mutant. We focused on a P_M_ inside *maf* gene that transcribes *radC*-*mreB*-*mreC*-*mreD*-*minC*-*minD* operon, a P_M_ upstream of *rodA* gene, and a P_M_ inside *murG* that contributes to transcription of the *murB*-*divIB*-*ylxW*-*ylxX*-*sbp*-*ftsA*-*ftsZ* operon (Fig 4E). Mutation of these three promoters individually or in combination had little effect on the colony size of the WT strain (Fig. S4E), since these genes are also expressed from σ^A^-dependent promoters. In a *yhdK* mutant, however, mutation of P_M_(*maf*) and P_M_ (*murG*) led to an additive increase of colony size, while mutation of P_M_(*rodA*) led to a small detrimental effect (Fig. 4F). We conclude that overexpression of genes downstream of P_M_(*maf*) and P_M_(*murG*) may overwhelm the membrane-protein insertion pathway, and thereby contribute to a net negative effect on cell fitness. Consistent with this idea, in cells expressing SpoIIIJ^Q140K^, the beneficial effect of mutating P_M_(*maf*) and P_M_(*murG*) was largely abolished (Fig. 4G).

## Discussion

The YidC membrane protein insertase is the bacterial representative of the YidC/Oxa1/Alb3 protein family of evolutionary conserved integral membrane proteins [18, 27]. In *E. coli*, YidC is required for the assembly of one or more essential protein complexes that reside in the inner membrane, including subunits of the energy generating F_1_F_O_ ATPase and NADH dehydrogenase I [34]. Although YidC can function in concert with the SecYEG translocon as a foldase/chaperone, YidC can also function independently for the insertion of small membrane proteins with one or two transmembrane segments [35].

*Bacillus subtilis* encodes two YidC paralogs, SpoIIIJ and YidC2 (formerly YqjG). Either protein can support growth of *B. subtilis*, but a double mutant is inviable. Under most conditions SpoIIIJ is the functional YidC homolog and is responsible for insertion of transmembrane proteins, including the F_1_F_O_ ATPase [19]. MifM, a membrane protein encoded upstream of *yidC2*, is also a substrate of SpoIIIJ and serves as a sensor for SpoIIIJ activity [21]. In the absence of SpoIIIJ activity, the translational pause caused by the membrane insertion of MifM is abolished and YidC2 is translationally activated [26, 36]. Although either SpoIIIJ or YidC2 can support growth, they appear to differ in the efficacy of inserting specific proteins: cells lacking SpoIIIJ are impaired in sporulation [20], whereas those lacking YidC2 have decreased competence [23]. The relationship between YidC protein sequence and the selection of specific client proteins is poorly understood. In cells depleted for both SpoIIIJ and YidC2 there was a substantial upregulation in expression of the Clp protease system and the LiaIH membrane-stress proteins that are regulated by the LiaRS two-component system [23]. This suggests that impairment of membrane protein insertion leads to an accumulation of misfolded proteins and disruption of the cell membrane.

One critical feature required for YidC function, as first visualized in the structure of the *B. halodurans* YidC protein [16], is a hydrophilic substrate-binding channel postulated to interact transiently with transmembrane segments of nascent integral membrane proteins. In YidC proteins the first transmembrane region (TM1) contains a conserved Arg residue that is essential for function in many, but not all, YidC orthologs [37]. This positively charged residue is postulated to form a salt-bridge with single-pass transmembrane client proteins with acidic residues in their amino-terminal region [16]. Remarkably, the function of this conserved Arg residues (R73 in *Bacillus* SpoIIIJ) can be replaced by Arg residues introduced at any of six other positions in transmembrane segments of YidC [25].

In this work, we have described an unusual variant of SpoIIIJ found in *B. subtilis* strain PY79 which contains the conserved R73 residue and additionally a second positively charged residue (K140) at a position that can functionally replace R73. This variant SpoIIIJ protein (SpoIIIJ^Q140K^) is necessary and sufficient for *B. subtilis* to survive high level expression of the σ^M^ regulon. By screening of a library of SpoIIIJ variants with different charges, we revealed that all SpoIIIJ proteins that enable cells to tolerate loss of *yhdL* contain at least two, and in some cases three, positively charged residues in this channel. We postulated that these SpoIIIJ variants may be more capable of accommodating the increased expression of membrane proteins under σ^M^ control. Support for this hypothesis derives from experiments in which the deletion of individual σ^M^-dependent promoters that control operons encoding multiple integral membrane proteins was found to improve fitness of strains with high σ^M^ activity. Thus we reason that the relevant feature of SpoIIIJ^Q140K^ is the presence of increased positive charge in the substrate-binding channel, which probably results in an increased insertase/foldase activity for specific σ^M^-regulated membrane proteins.

Our results have clarified the nature of the lethality associated with a *yhdL* deletion that unleashes high level σ^M^ activity. In strains with elevated σ^M^ activity there is an increased flux of proteins targeted to the inner membrane and a subset of these may be inefficiently inserted by the native SpoIIIJ protein. This can result in a jamming of YidC-dependent protein translocation and a disruption of membrane function. The downstream sequelae associated with this disruption includes misfolding of proteins and induction of the secretion stress response (CssR), as well as genes associated with oxidative stress and DNA damage. These types of stresses may explain, in part, the inclusion of appropriate compensatory functions (including Spx, some DNA repair functions, and SecDF) as part of the σ^M^ regulon. It is presently unclear why the *spoIIIJ^Q140K^* allele arose in the PY79 strain, or what conditions may have led to its selection. However, this strain was derived from strains treated with chemical mutagens to cure the endogenous prophage SPß [14] and this or other selection conditions may have contributed to emergence of this mutation. Further studies will be required to better understand how variations in YidC structure can fine-tune the substrate selectivity of this essential membrane protein insertase, and the stresses that arise when this system is challenged by the induction of highly expressed membrane proteins.

## Material and Methods

### Strains, plasmids and growth condition

All strains used in this work are listed in Table S5, and all DNA primers are listed in Table S6. Bacteria were routinely grown in liquid lysogeny broth (LB) with vigorous shaking, or on plates (1.5% agar; Difco) at 37 °C unless otherwise stated. LB medium contains 10 g tryptone, 5 g yeast extract, and 5 g NaCl per liter. Plasmids were constructed using standard methods [38], and amplified in *E. coli* DH5α or TG1 before transforming into *B. subtilis*. For selection of transformants, 100 μg ml^-1^ ampicillin or 30 μg ml^-1^ kanamycin was used for *E. coli*. Antibiotics used for selection of *B. subtilis* transformants include: kanamycin 15 μg ml^-1^, spectinomycin 100 μg ml^-1^, macrolide-lincosamide-streptogramin B (MLS, contains 1 μg ml^-1^ erythromycin and 25 μg ml^-1^ lincomycin), and chloramphenicol 10 μg ml^-1^. For spot dilution assays, cells were first grown in liquid culture at 37 °C with shaking to mid-exponential phase (OD_600_∼0.3-0.4), washed twice in LB medium without inducer, then serial diluted in LB medium without inducer. 10 μl of each diluted culture was then spotted onto plates and allowed to dry before incubation at 37 °C for 12-24 hours.

### Genetic techniques

Chromosomal and plasmid DNA transformation was performed as previously stated [39]. The pPL82 plasmid-based P_spac(hy)_ overexpression constructs were sequencing confirmed before linearized and integrated into the *amyE* locus [24]. The pAX01 plasmid-based P*_xylA_* overexpression constructs were sequencing confirmed before linearized and integrated into the *ganA* locus [40]. The *ganA*::P*_xylA_*-*yhdL*-*cat* and *thrC*::P_M_-*spoVG*-*lacZ*-*spec* constructs were made with LFH PCR to avoid antibiotic marker conflicts [10]. Markerless in-frame deletion mutants were constructed from BKE or BKK strains as described [41]. Briefly, BKE or BKK strains were acquired from the *Bacillus* Genetics Stock Center (http://www.bgsc.org), chromosomal DNA was extracted, and the mutation containing an *erm*^R^ (for BKE strains) or *kan*^R^ (for BKK strains) cassette was transformed into our WT 168 strain. The antibiotic cassette was subsequently removed by introduction of the Cre recombinase carried on plasmid pDR244, which was later cured by growing at the non-permissive temperature of 42 °C. Gene deletions were confirmed by PCR screening using flanking primers. Unless otherwise described, all PCR products were generated using *B. subtilis* 168 strain chromosomal DNA as template. DNA fragments used for gene over-expression were verified by sequencing. Null mutant constructions were verified by PCR.

Mutations to selectively inactivate σ^M^-dependent promoters were generated by either promoter deletion (*rodA*) or by inactivating point mutations (*murG*, *maf*, *spx*). To inactivate P_M_(*rodA*), a 91 bp region containing the P_M_ (located upstream of a σ^A^ -dependent promoter) was deleted using CRISPR. The resulting deletion had a junction sequence of: CACATTATCGC/TTTCGTGTAGC. The point mutations inactivating *murG* and *maf* both changed the −10 region sequence from consensus, CGTC, to TGTT. The P_M_* mutation inactivating the promoter regulating the *yjbC-spx* operon is a 3 bp substitution changing the −10 region from consensus, CGTC, to AAGT, as previously described [42].

Mutations of *spoIIIJ* at native locus, as well as ΔP_M_-*rodA* were constructed using a <u>c</u>lustered <u>r</u>egularly interspaced <u>s</u>hort <u>p</u>alindromic repeats (CRISPR)-based mutagenesis method as previously described [10]. Briefly, possible protospacer adjacent motifs (PAM) site, which is NGG for *Streptococcus pyogenes* Cas9, was identified and off-target sites were checked against *B. subtilis* genome using BLAST. If no off-target site was identified, the PAM site was chosen and 20 bps upstream of the site were used as sgRNA and cloned into vector pJOE8999 [43]. The repair template was generated by joining two or more PCR products, with intended mutation (with additional mutation to abolish gRNA recognition if necessary) introduced by PCR primers, and cloned into the pJOE8999-sgRNA vector. DNA sequence for amino acid substitution was chosen according to the preferred codons of *B. subtilis* [44]. The pJOE8999 derivative containing both the sgRNA and repair template was then cloned into competent cells of *E. coli* strain TG1 to produce concatemer plasmids, which were transformed into *B. subtilis* at 30°C. Transformants were then grown at 42°C to cure the plasmid, and intended mutations were confirmed by sequencing. ΔP_M_-*rodA* was constructed with repair template amplified using primers 7426, 7427, 7428 and 7429, and the gRNA constructed using 7430 and 7431. SpoIIIJ^Q140K^ mutation in 168 was constructed with repair template amplified using primers 7868, 7869, 7870 and 7871, and the gRNA constructed using 7866 and 7867. SpoIIIJ^K140Q^ mutation in PY79 was constructed with repair template amplified using primers 7866, 8247, 8246 and 7871, and the gRNA constructed using 8248 and 8249. SpoIIIJ^R73A^ mutation was constructed with repair template amplified using strain 168 genomic DNA as template with primers 7868, 8280, 8281 and 7871, and the gRNA constructed using 8278 and 8279. SpoIIIJ^R73AQ140K^ mutation was constructed with the same primer sets for SpoIIIJ^R73A^ mutation, expect the repair template was amplified using strain PY79 genomic DNA. All constructs were sequencing confirmed.

SpoIIIJ library of different positive charges were constructed using degenerate primers and LFH PCR. Four DNA fragments were amplified and joined using LFH PCR. The fragments include one with part of gene *hom* and *thrC* (amplified with primers 8766 and 8767), one with the spoIIIJ gene from 168 (amplified with primers 8768 and 8681, containing the native promoter and ribosomal binding site), one with a spec^R^ cassette (amplified with primers 8682 and 8769), and one with part of *thrC* and full length of *thrB* (amplified with primers 8770 and 8771). The joined PCR product was first transformed into strain 168 to generate HB23976. Then using genomic DNA of HB23976 as template, four new DNA fragments were amplified using degenerate primer pairs 8766 and 8689, 8686 and 8690, 8687 and 8691, and 8688 and 8771. These fragments were joined together using LFH PCR and the joined PCR product (more than 10 μg DNA) was expected to contain all 128 possible combinations of SpoIIIJ charge from 0 to +7. The PCR product library was used to transform strain 168, and more than ten-thousand transformants (from 20 plates, each contains more than 500 colonies) were pooled together to extract genomic DNA to form a virtually equivalent DNA library of the *spoIIIJ* variants. This genomic DNA library provides high transformation efficiency and was used to transform a *yhdL* depletion strain (HB23953) and transformants were selected on LB plate supplemented with X-gal but no xylose for *yhdL* induction. More than 200 transformants were re-streaked onto fresh LB plate with X-gal but no xylose to confirm robust growth under high SigM condition, and 103 of them were Sanger sequenced for the *thrC*-*spoIIIJ*-*spec* region to identify the *spoIIIJ* variants.

### Whole genome re-sequencing and sequence analysis

Chromosomal DNA of suppressor strains was extracted using Qiagen DNeasy Blood & Tissue Kit. DNA was then sent to Cornell University Institute of Biotechnology for sequencing using Illumina HiSeq2500 with Single-end 100 bp reads. Sequencing results were analyzed using CLC workbench version 8.5.1 and mapped to the genome of strain 168 (reference accession number NC_000964.3). Note that our working stock of *B. subtilis* 168 has 21 SNPs compared to the cited reference sequence, and these common SNPs were not considered, and only newly introduced SNPs from the PY79 strain were tabulated (Table S2). Unmapped reads were de novo assembled and contigs larger than 1 kb were BLASTed against genome of strain PY79 (reference accession number NC_022898.1). Single nucleotide variants (SNVs) were detected using default settings, and gene deletions larger than 300 bps were identified by manually scanning regions of low coverage.

### Colony size measurement

Colony size was measured using Fiji Image J [45]. Briefly, bacterial cells were grown in liquid LB medium at 37°C with vigorous shaking to mid-exponential phase (OD_600_∼0.3-0.4), then serial diluted to desired concentrations. Diluted cells were plated onto fresh LB plates (15 ml medium per plate, the diameter of the plate is 10 cm and the height 15 mm, VWR, US, Catalog number 25384-342), and multiple dilutions were used. Plates were incubated at 37 °C for 24 hours. Plates containing less than 100 separate single colonies were used for size measurement, because this number of colonies per plate ensures sufficient sample size and does not cause reduced colony size due to crowdedness and nutrient limitation. Pictures of plates were taken with a ruler as a length reference, and colony size was measured using Fiji Image J per software’s instruction. For each strain, at least 100 colonies were measured, and box and whisker plots were used.

### Luciferase reporter construction and measurement

Luciferase reporter construction and measurement was performed as previous described [46]. The luciferase reporters were constructed by inserting the tested promoters into the multicloning sites of pBS3C*lux* [47]. The promoter P*_htrA_* was amplified using primers 8403 and 8404. For luciferase measurements, 1 μl of exponentially growing cells were inoculated into 99 μl of fresh medium in a 96 well plate, incubated at 37 °C with shaking using a SpectraMax i3x plate reader, and OD_600_ and luminescence were measured every 12 min. The data was analyzed using SoftMax Pro 7.0 software. Promoter activity was normalized by dividing the relative light units (RLU) by OD_600_.

### Phase contrast microscopy

Cells were grown in liquid LB medium to exponential phase (OD_600_ ∼0.4) and loaded on saline (0.90% NaCl, w/v) agarose pads (0.8% final concentration) on a glass slide. Phase contrast images were taken using a Leica DMi8 microscope equipped with a 100x immersion objective and Leica Application Suite X software.

### Western blot

Western blot was performed as described previously [42]. Briefly, cells were grown in 5 ml LB medium in a 20 ml test tube a 37 °C with vigorous shaking. Inducer IPTG or xylose were added when required by the construct to induce SpoIIIJ. After reaching exponential phase (OD_600_∼0.3-0.4), 1 ml cells were pelleted by centrifugation at 4°C, resuspended in 100 μl pre-chilled buffer (containing 25 μl 4X Laemmli Sample buffer (Bio-Rad, USA), 10 μl 1M DTT, 65 μl H_2_O) and kept on ice. Cells were then lysed and crude cell lysate was loaded to a 4-20% SDS-PAGE stain-free gel for electrophoresis. The gel was visualized using ImageLab with stain-free gel protocol. Proteins were then transferred onto a PVDF membrane using the TransBlot Turbo Transfer System (Bio-Rad, USA), and immune blotting was performed using anti-SpoIIIJ antiserum. The blot was visualized using the Clarity Western ECL substrate (Bio-Rad) and ImageLab software. Band intensity was calculated using the ImageLab software and normalized using total protein amount according to SDS-PAGE gel image.

### β-galactosidase assay for *yidC2’*-*lacZ*

Bacterial cells containing the *yidC2’*-*lacZ* translational fusion were grown to late exponential phase (OD_600_ 0.6-0.8) in 96-well plates with 200 μl LB medium per well at 37 °C with vigorous shaking. Cells were pelleted by centrifugation, resuspended in Z buffer supplemented with DTT (Dithiothreitol, 400 nM final concentration), and lysed by lysozyme. OD_600_ was measured before lysozyme treatment. After lysis, ONPG (ortho-nitrophenyl-β-galactoside) was added and OD_420_ and OD_550_ were measured every 2 minutes. Product accumulation was calculated using formula product=1000×[OD_420_-(1.75×OD_550_)] and plotted against time. The slope of the linear part of the product accumulation curve was calculated using Excel and Miller Unit (MU) was calculated using formula MU=Slope/OD_600_/V, where V is the volume of cells used for the reaction (200 μl).

### RNA extraction and qRT-PCR

Cells were grown to mid-exponential phase (OD_600_ ∼0.4) and RNA was extracted using RNeasy Mini Kit (Qiagen). The extracted was then treated with DNase I (Invitrogen) and the quality of RNA was checked with electrophoresis. RNA was then reverse transcribed into cDNA using High-Capacity cDNA Reverse Transcription Kit (Applied Biosystems). Quantitative real-time PCR (qRT-PCR) was performed using SYBR Green (BioRad) and the *topA* gene was used for reference of data normalization.

## **Acknowledgements**

We thank Dr. Shinobu Chiba for providing the *yidC2’*-*lacZ* translational fusion reporter and antiserum for SpoIIIJ, Dr. David Rudner and Alexander Meeske for strain BAM1077, Dr. Vaidehi Patel and Wen-Wen Zhou for providing the P_M_* mutations for the *murG* and *maf* operons, and Dr. Tobias Doerr for valuable discussions and sharing equipment. We also thank Dr. Pete Chandrangsu and Ahmed Gaballa for help with experimental design and interpretation, and Daniel Roistacher for helping with experiments.

## Author Contributions

Conceptualization: Heng Zhao, John D. Helmann

Funding acquisition: John D. Helmann.

Investigation: Heng Zhao, Anita Sachla

Project administration: John D. Helmann.

Writing - original draft: Heng Zhao, John D. Helmann.

Writing - review & editing: Heng Zhao, Anita Sachla, John D. Helmann.

## Supplemental Figure legends

**Fig. S1.**
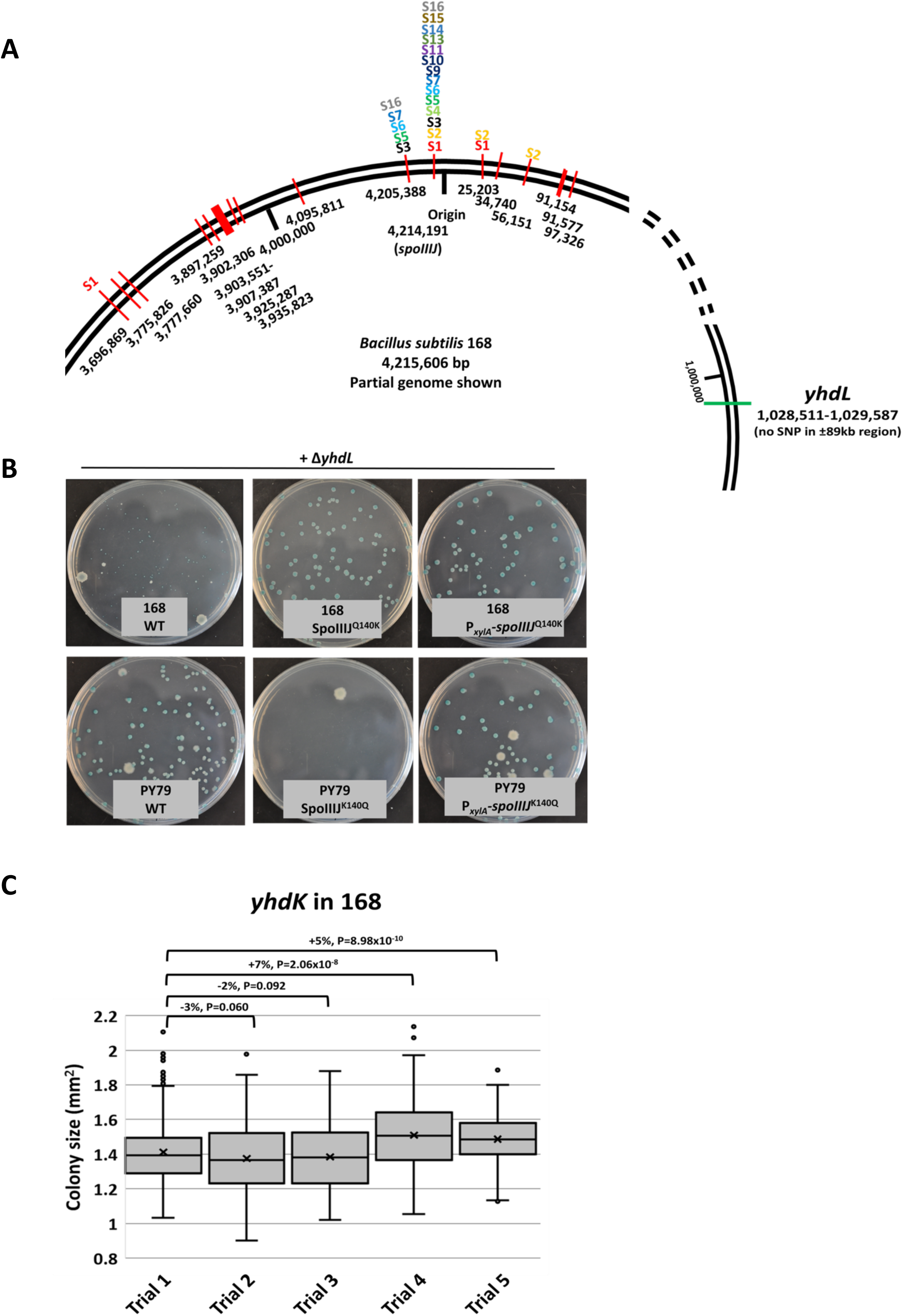
**A**. Map of SNPs from strain PY79 to 168 and distribution of SNPs contained in each congression suppressors. Genome coordinates were based on the 168 reference genome with NCBI accession number NC_000964.3. **B**. Transformation plates of *yhdL*::*kan* allele transformed into different strain backgrounds, selected on LB plates supplemented with kanamycin, X-gal and 1% xylose. **C**. Variation of *yhdK* colony size measurement between trials on different days with different batches of LB plates. P value was calculated using Student’s t test, and percentage changes of average colony size were shown.

**Fig. S2.**
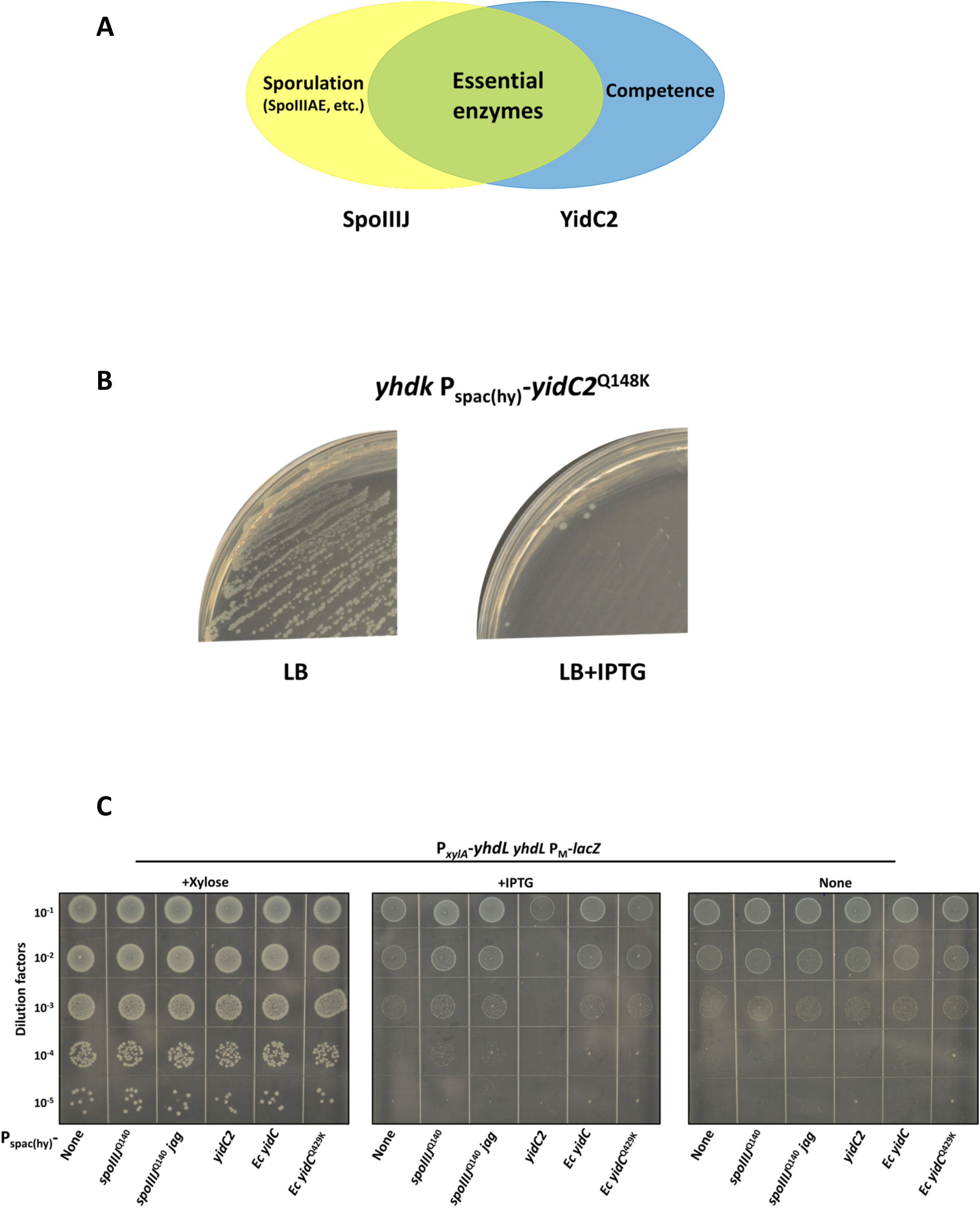
**A**. Venn diagram of function overlap and distinction of SpoIIIJ and YidC2 of *Bacillus subtilis*. **B**. Streaking of *yhdK* P_spac(hy)_-*yidC2*^Q148K^ on plates of LB or LB supplemented with 1 mM IPTG (final concentration). **C**. Spot dilution of *yhdL* depletion strains with P_spac(hy)_ based overexpression of different YidC homologs, on LB plates supplemented with a final concentration of 1% xylose (+Xylose), 1 mM IPTG (+IPTG) or nothing (None).

**Fig. S3.**
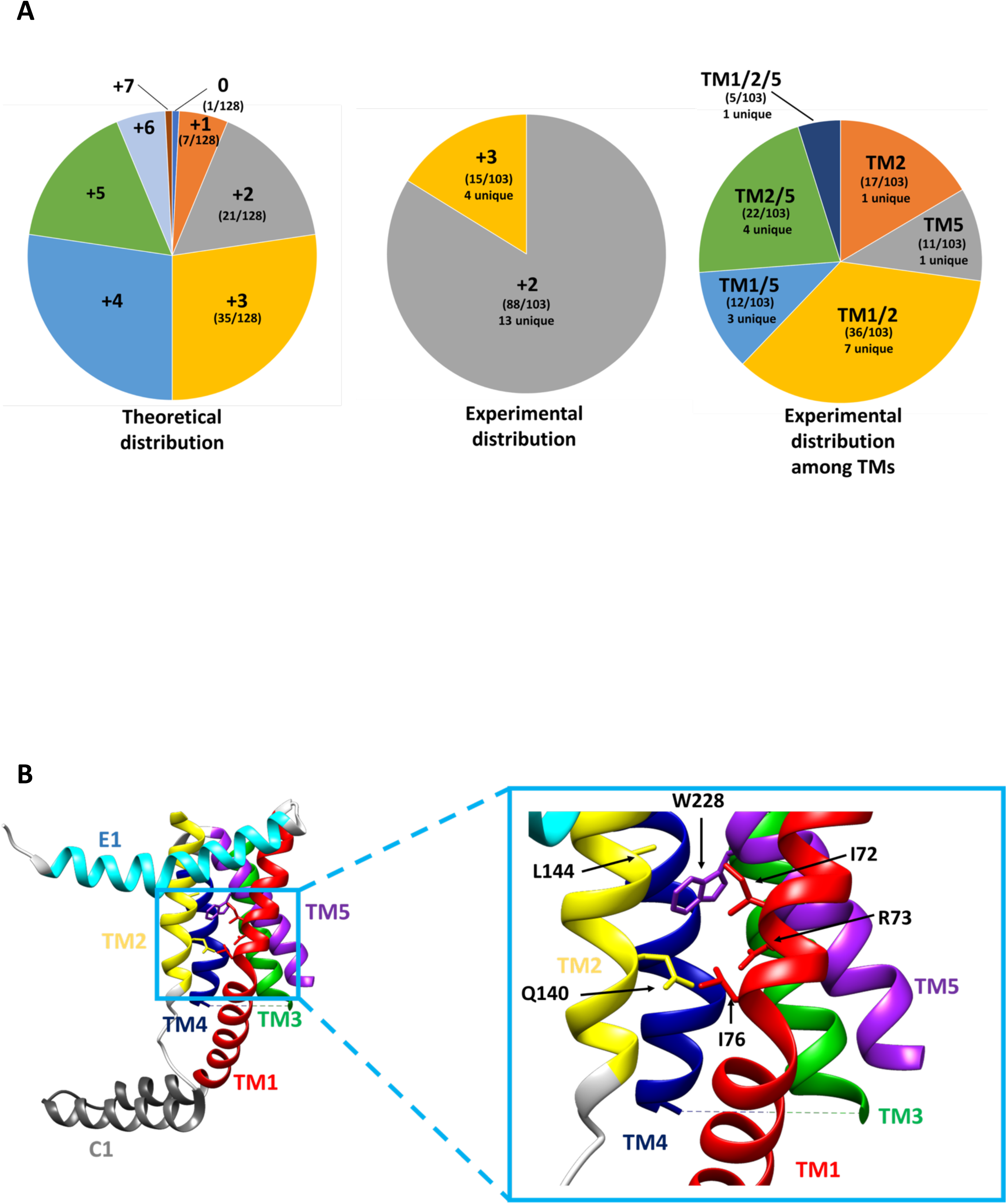
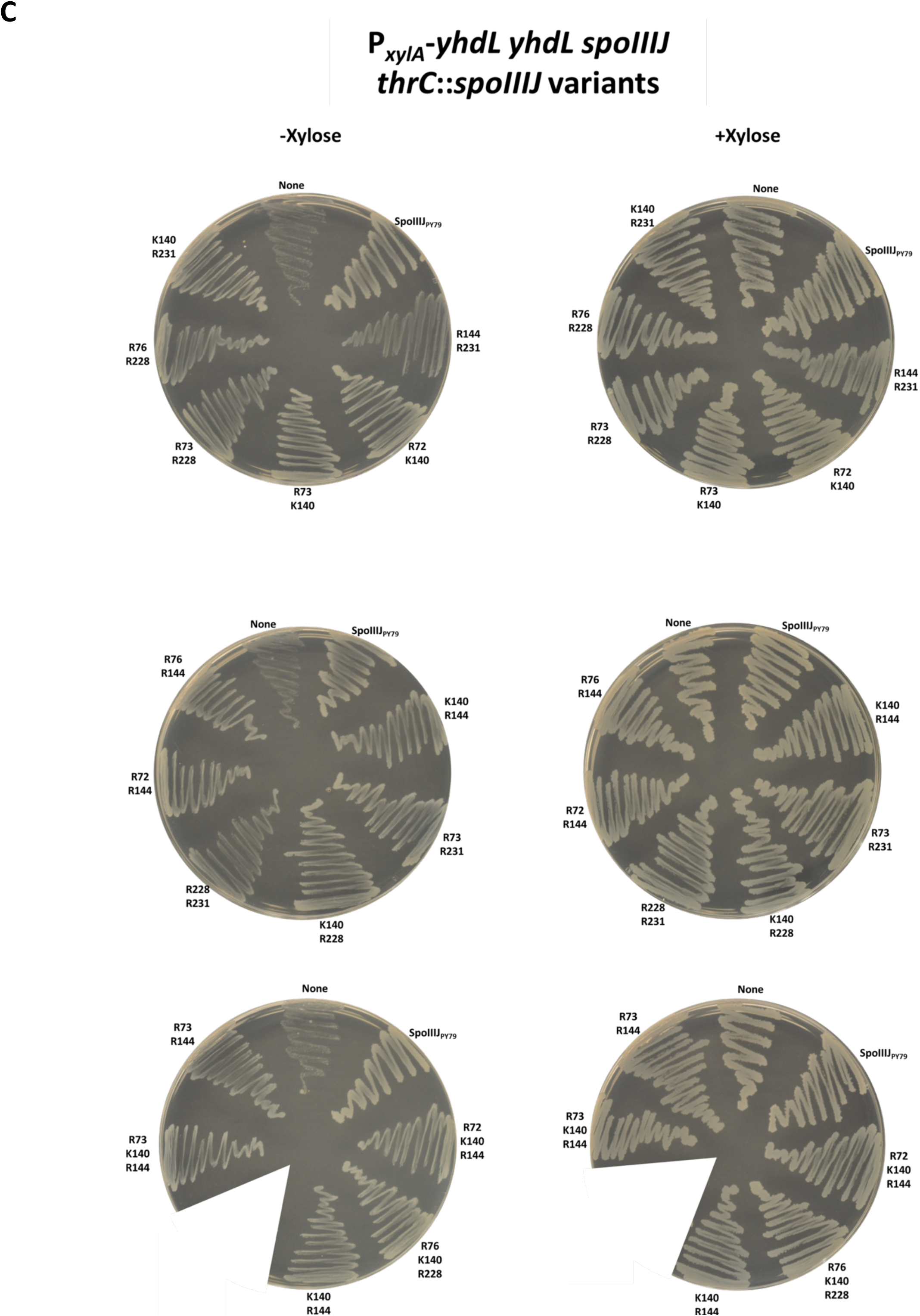
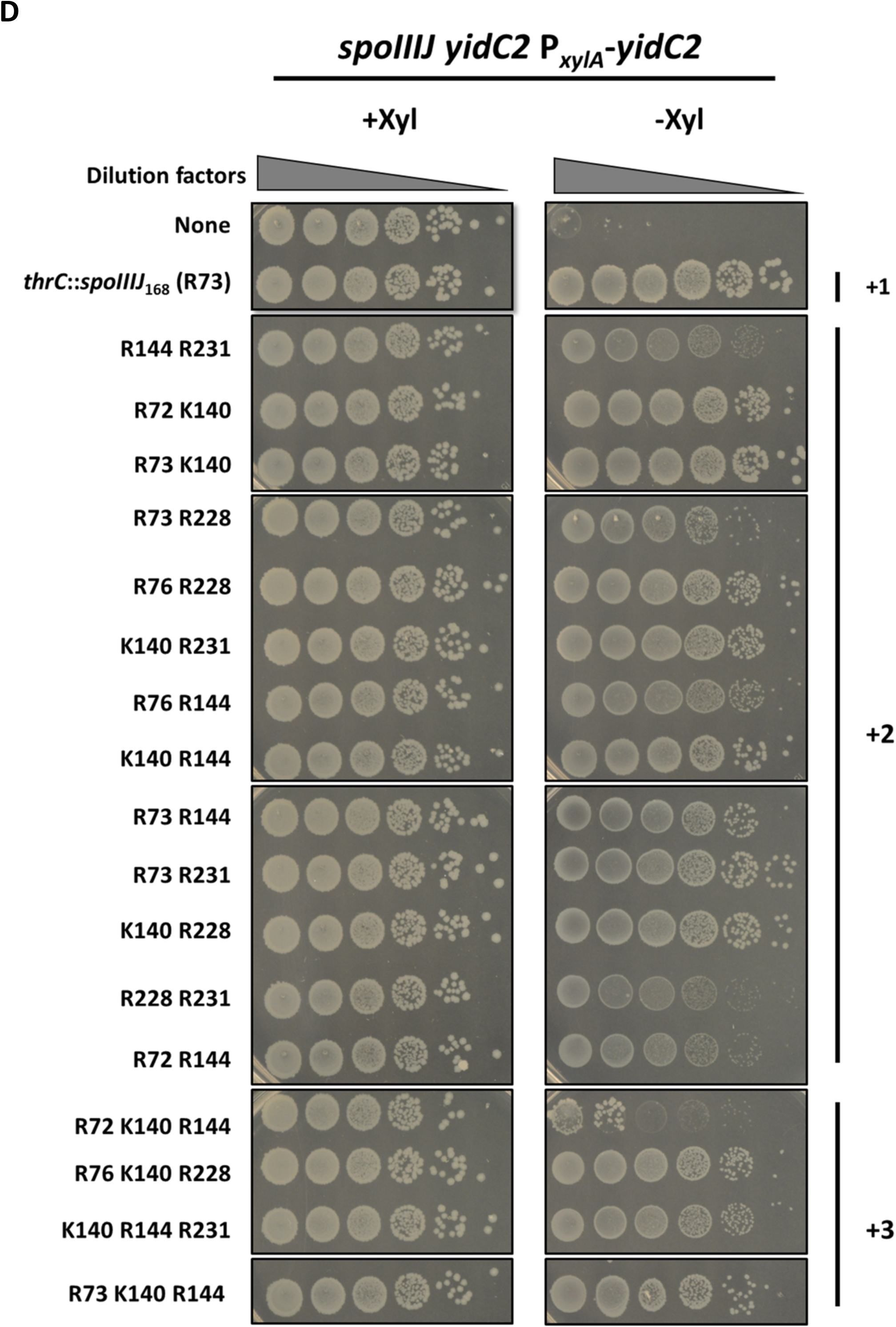
**A**. Distribution of 128 possible charges of SpoIIIJ in the *spoIIIJ* variants library. Theoretically, the majority of SpoIIIJ in the input DNA library contains +2 to +5 charges, while experimental data from 103 samples suggests the ones capable of providing high SigM tolerance contain +2 or +3 charges. Among the 103 samples with +2 or +3 charges, the positive charge can be located in one, two or three transmembrane segments, with the exception that no sample contains more than one positive charge in TM1 alone. **B**. Crystal structure of YidC from *Bacillus halodurans* (PDB ID 3WO6), showing the seven variable amino acid providing positive charges in the hydrophilic substrate binding chamber of the enzyme. Gly231 is not visible due to the lack of side chain of this residue. TM1-5, transmembrane region 1-5; E1, extracytoplasmic region 1; C1, cytoplasmic region 1.This figure was generated using UCSF Chimera 1.13[48]. **C**. Streaking of *yhdL* depletion strains with SpoIIIJ variants on LB plates with or without xylose inducer for *yhdL*. The positive charge of each variant was labelled next to the streaking, with a negative control “None” meaning no *spoIIIJ* variants at *thrC* locus (weak growth due to the depletion conditions, and cannot be restreaked), and a positive control “SpoIIIJ_PY79_” meaning the native SpoIIIJ mutated into the PY79 Q140K version (HB23719). **D**. Spot dilution of YidC depletion strains with *spoIIIJ* variants at *thrC* locus. The depletion strain has its native *spoIIIJ* and *yidC2* deleted, and a xylose inducible copy of P*_xylA_*-*yidC2*. Diluted cultures were spotted on LB without xylose (-Xyl) or with 1% final concentration of xylose (+Xyl). The negative control (None) has no *spoIIIJ* at *thrC* locus, while the positive control has the 168 version of *spoIIIJ* that contains a single positive charge at R73. Some SpoIIIJ variants exhibited reduced growth ability, and the variant containing R72 K140 R144 failed to growth, although the emergence of suppressors was noted.

**Fig. S4.**
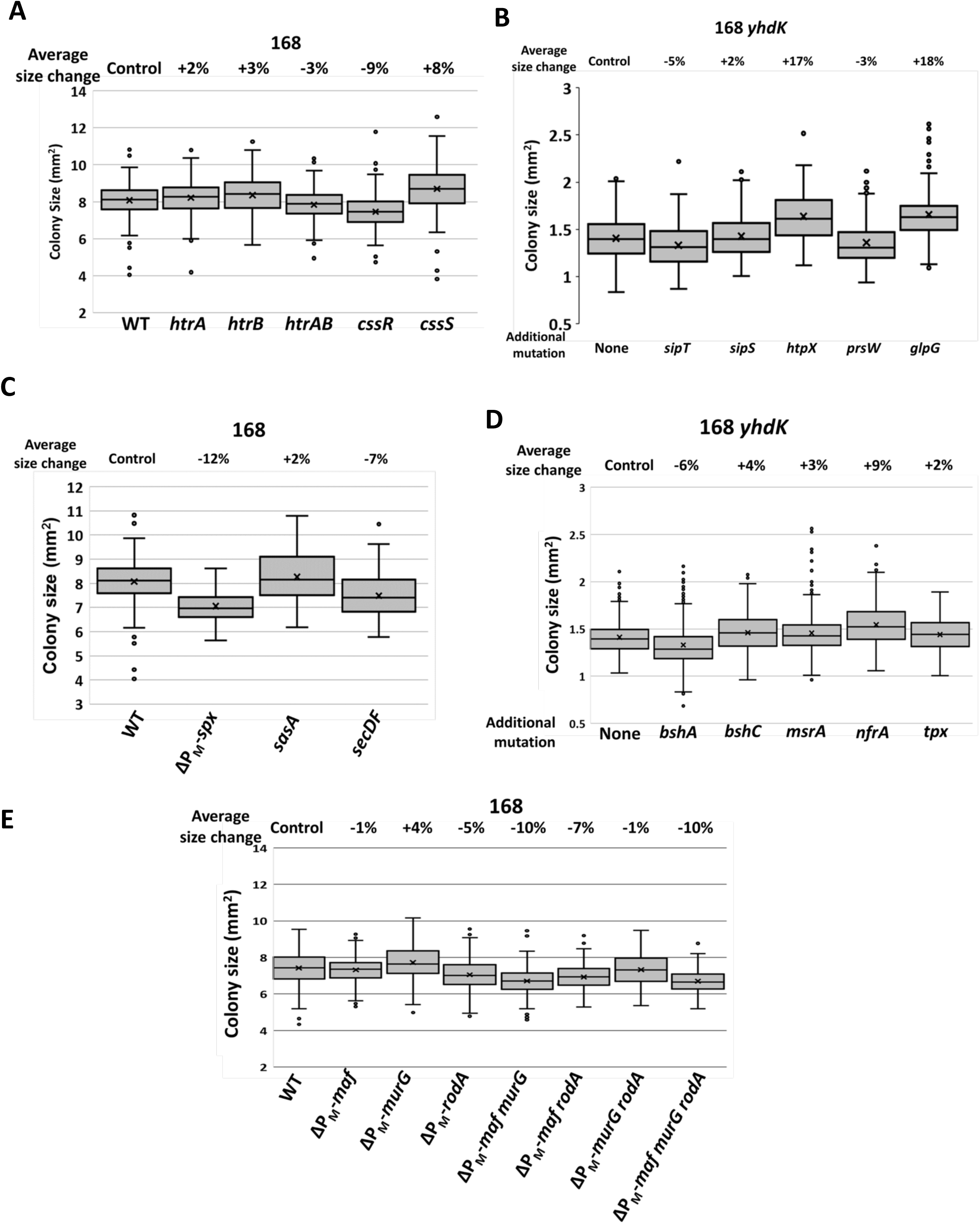
Colony size of different strains. Percentage changes of average colony size were shown on each Box and Whisker plot.

**Table S1.**
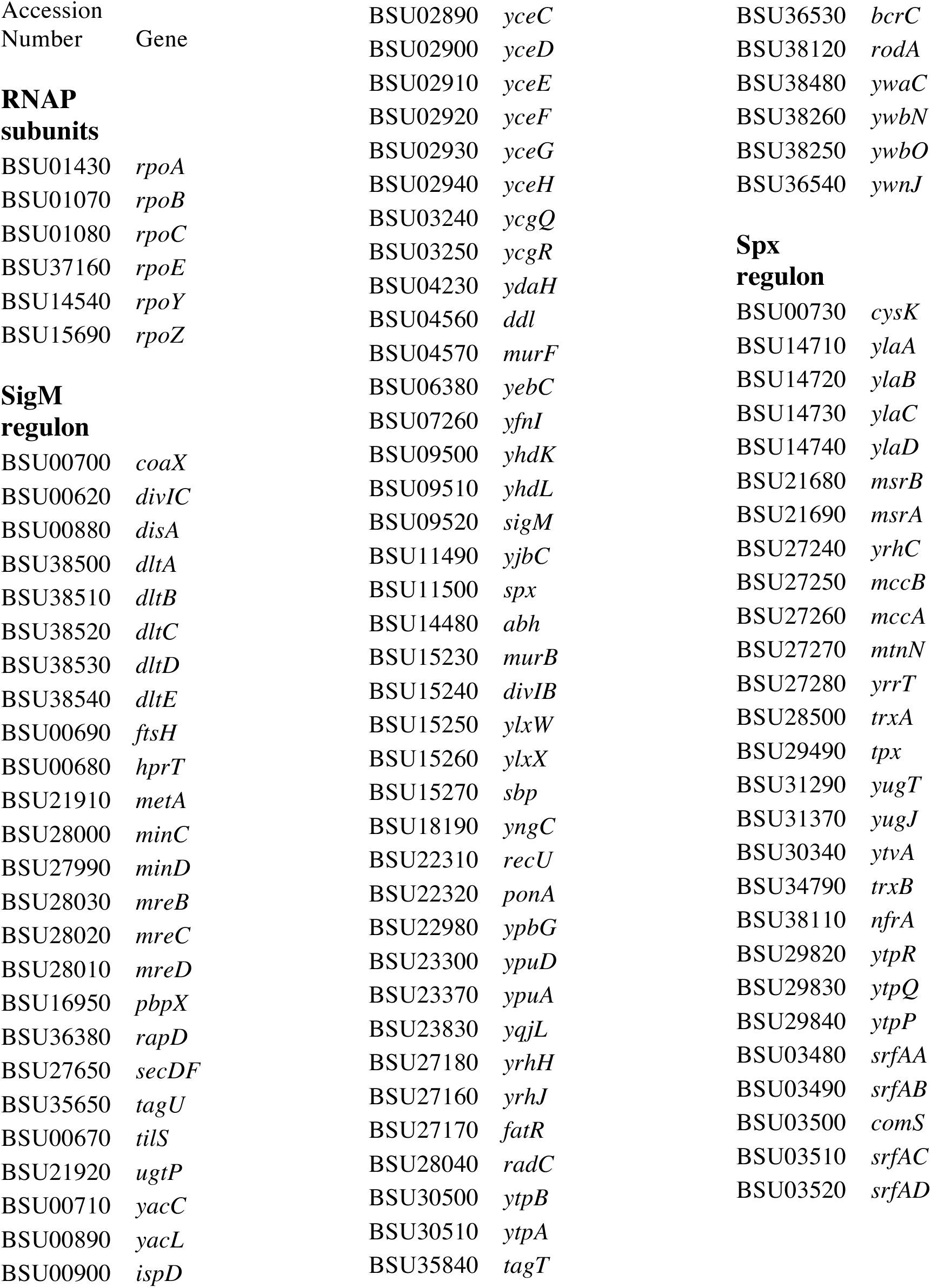
Genes compared between reference genomes of *B. subtilis* 168 (NCBI accession number NC_000964.3) and PY79 (NC_022898.1). No mutations were found in these genes.

**Table S2.**
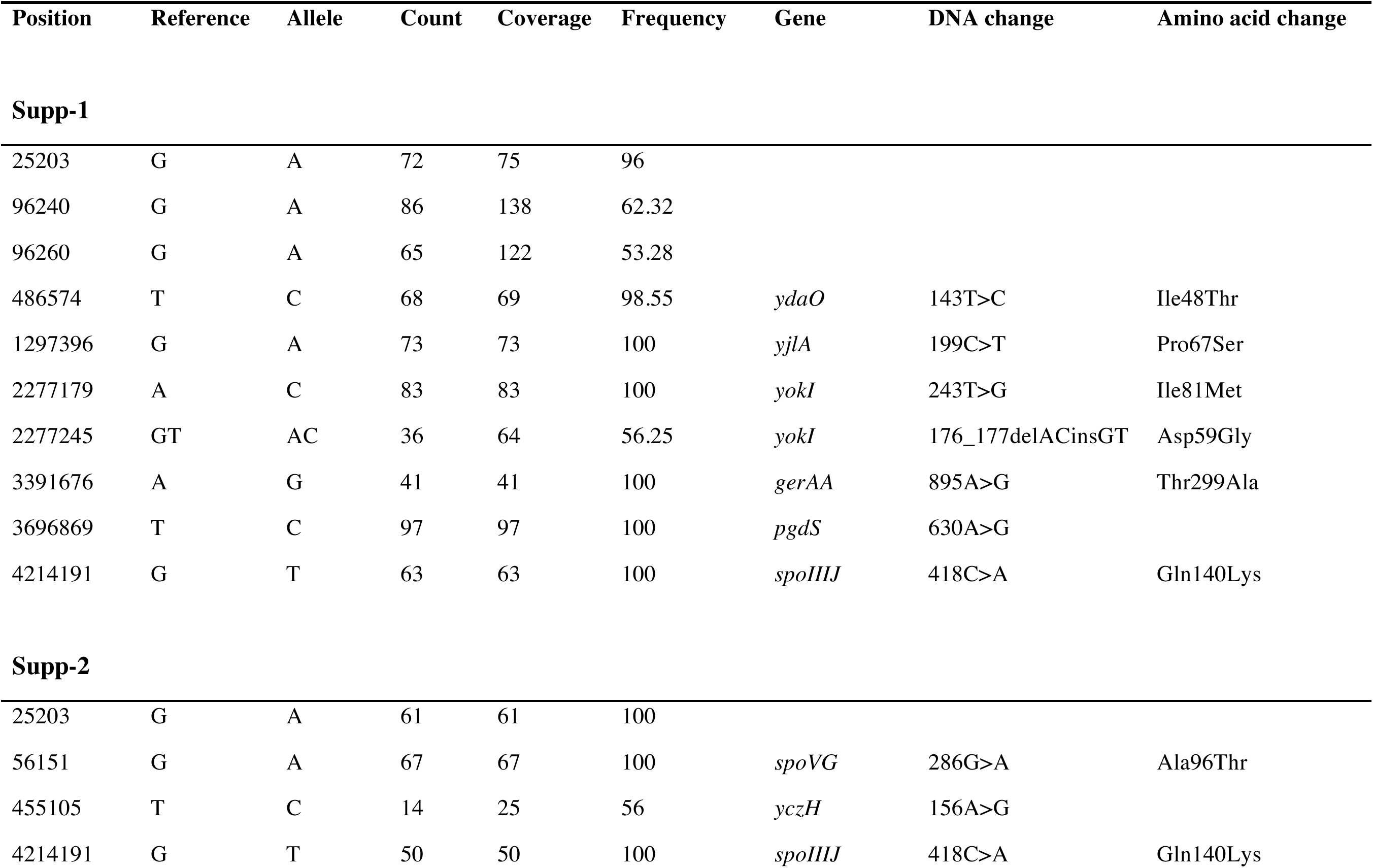
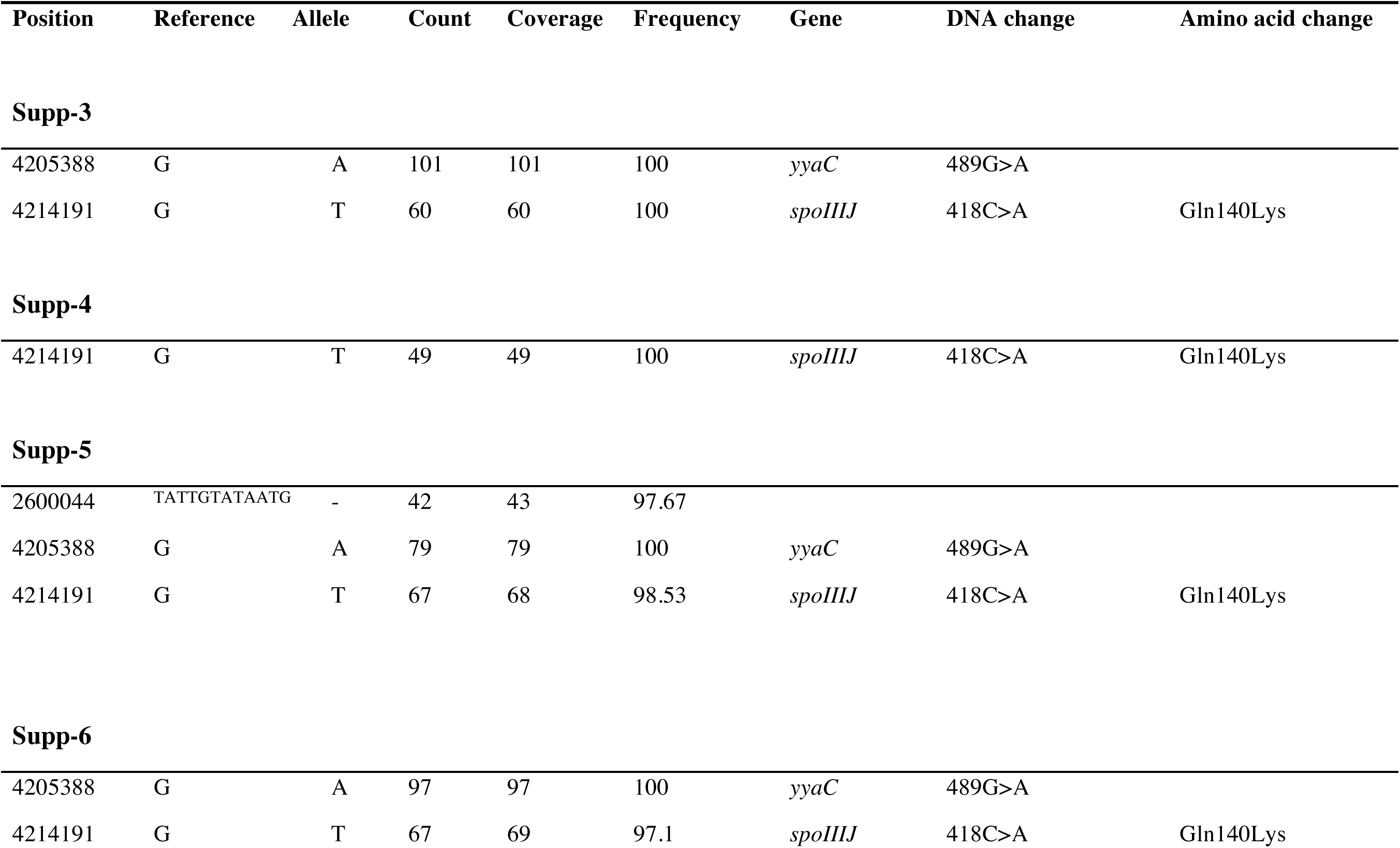
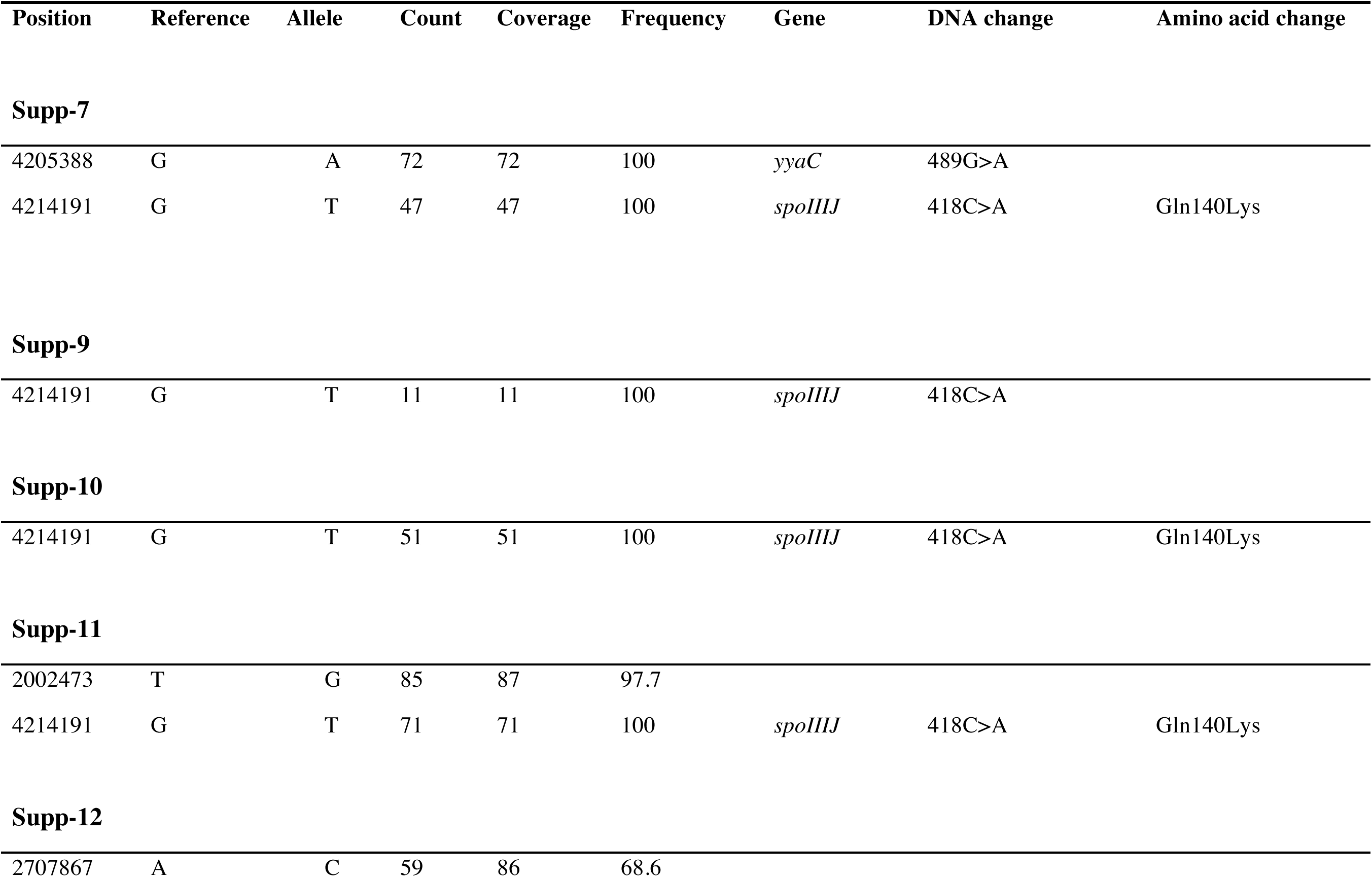
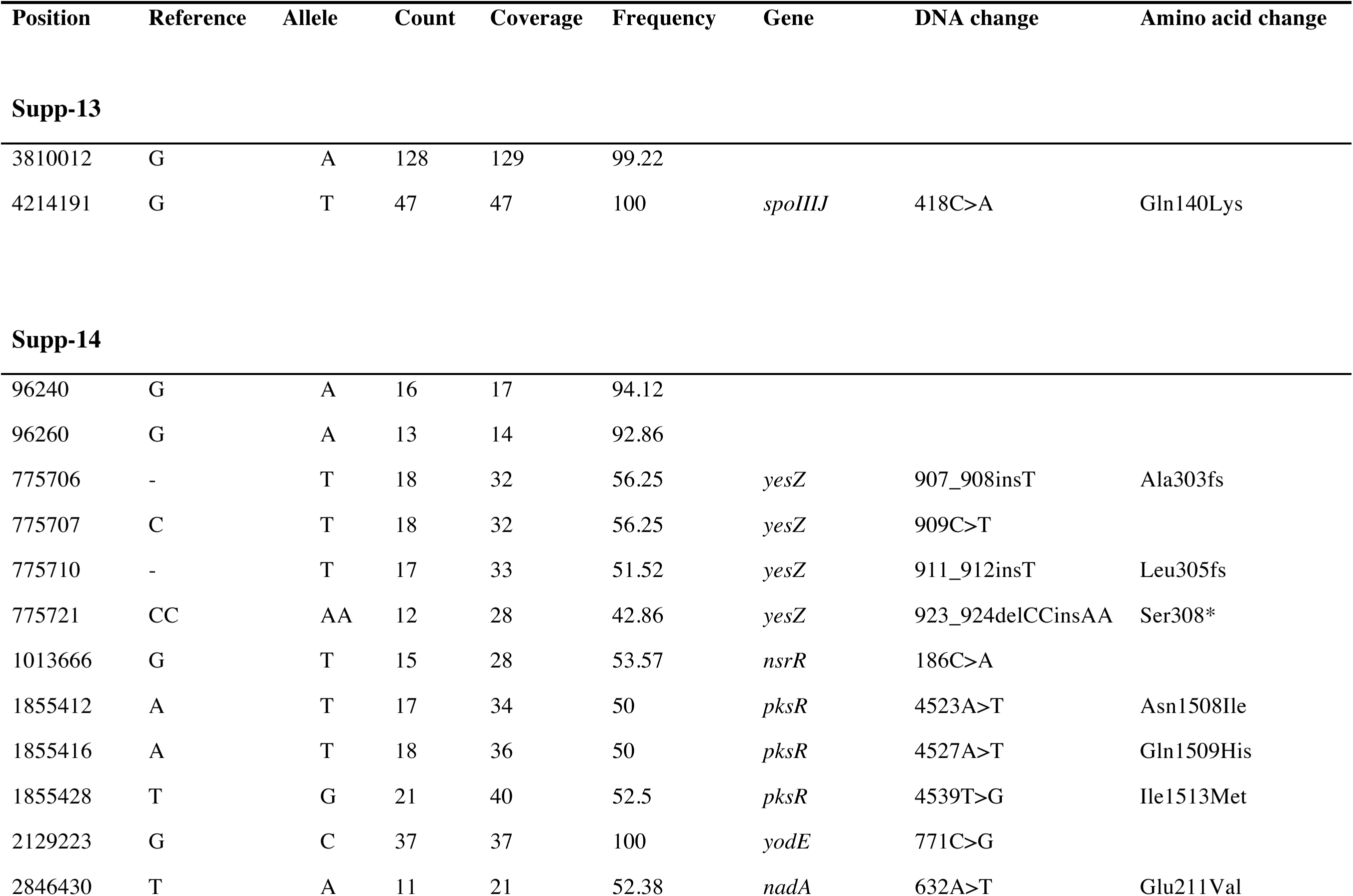
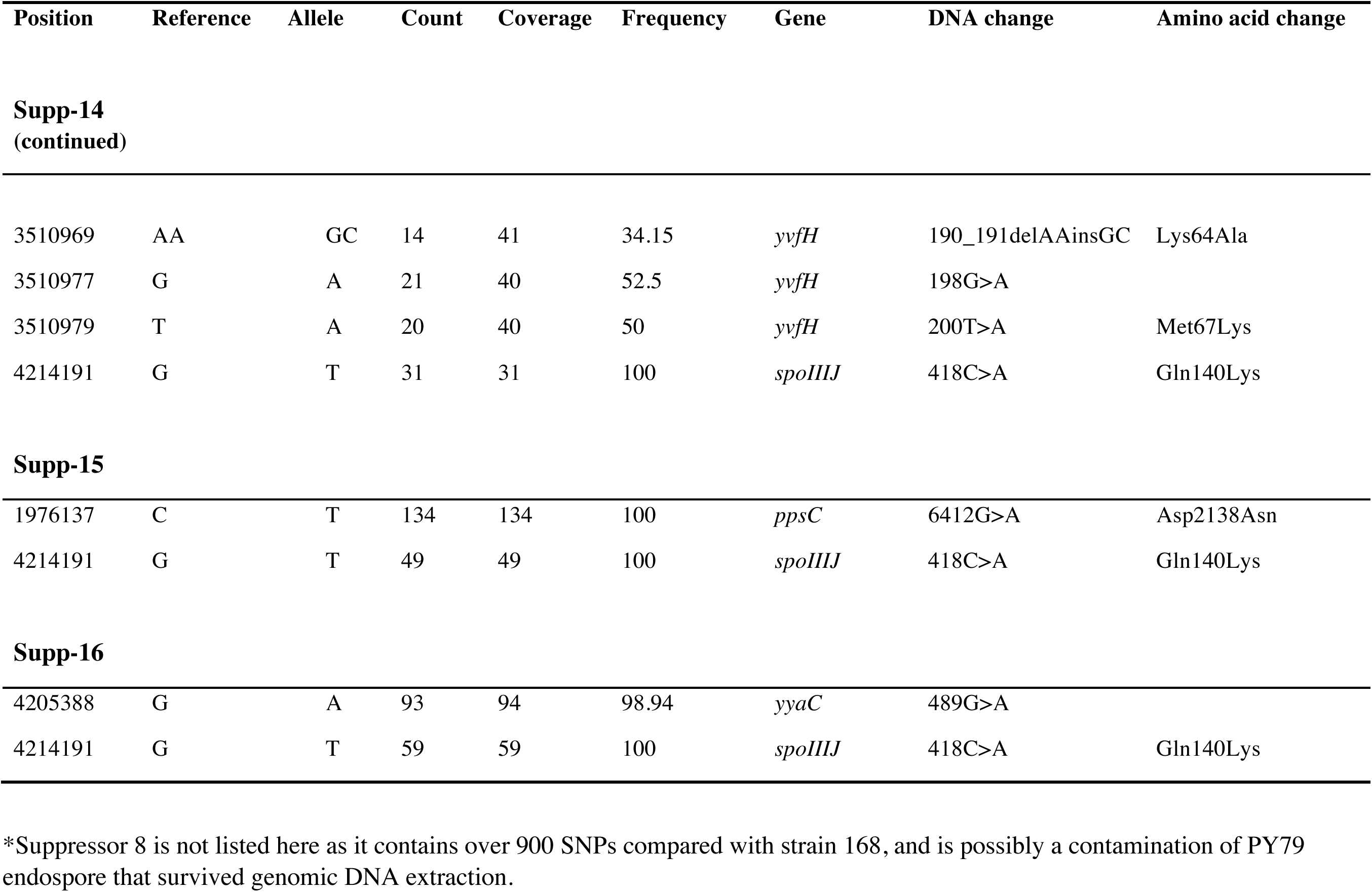
Whole genome sequencing of transformants generated by congression.

**Table S3.**
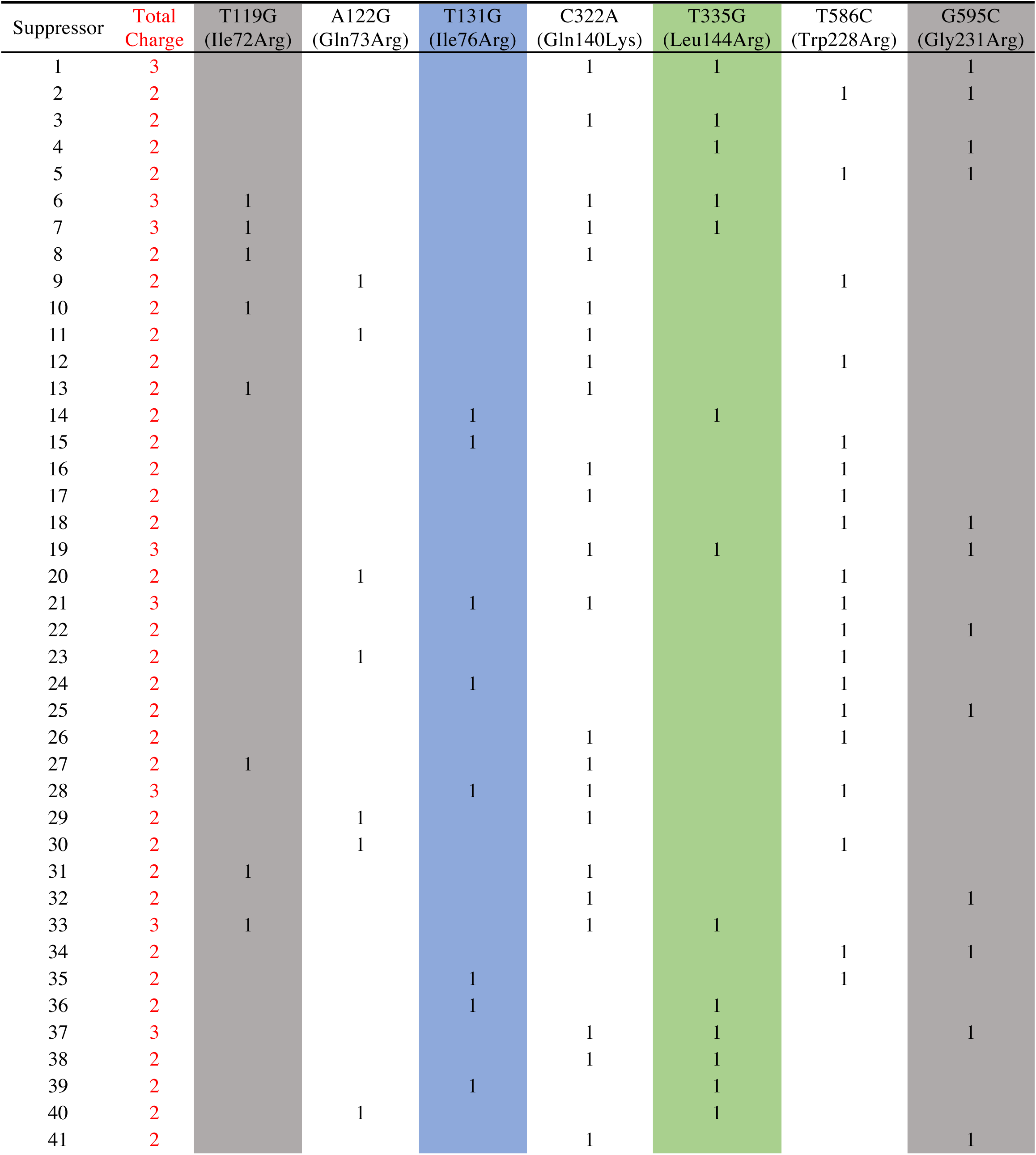
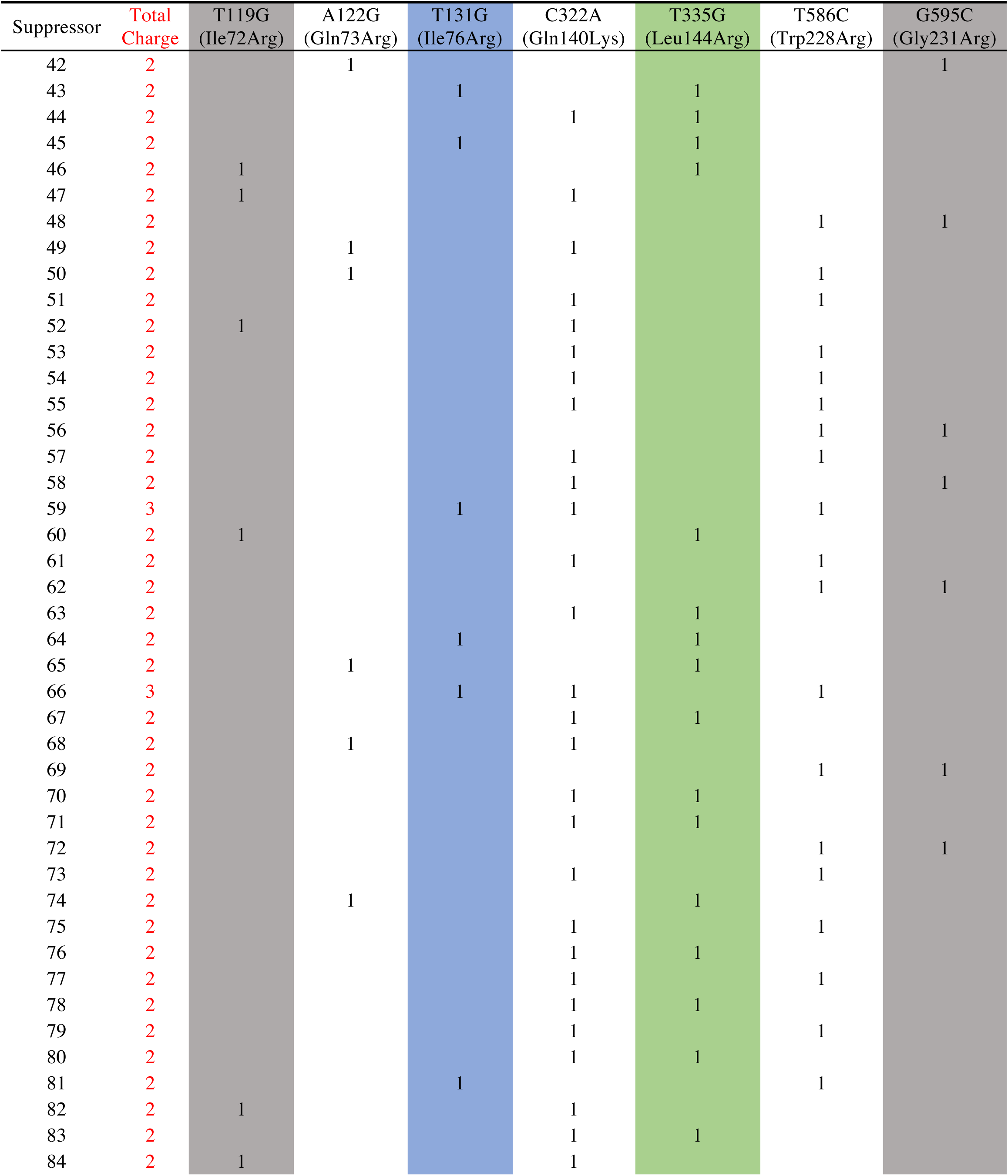
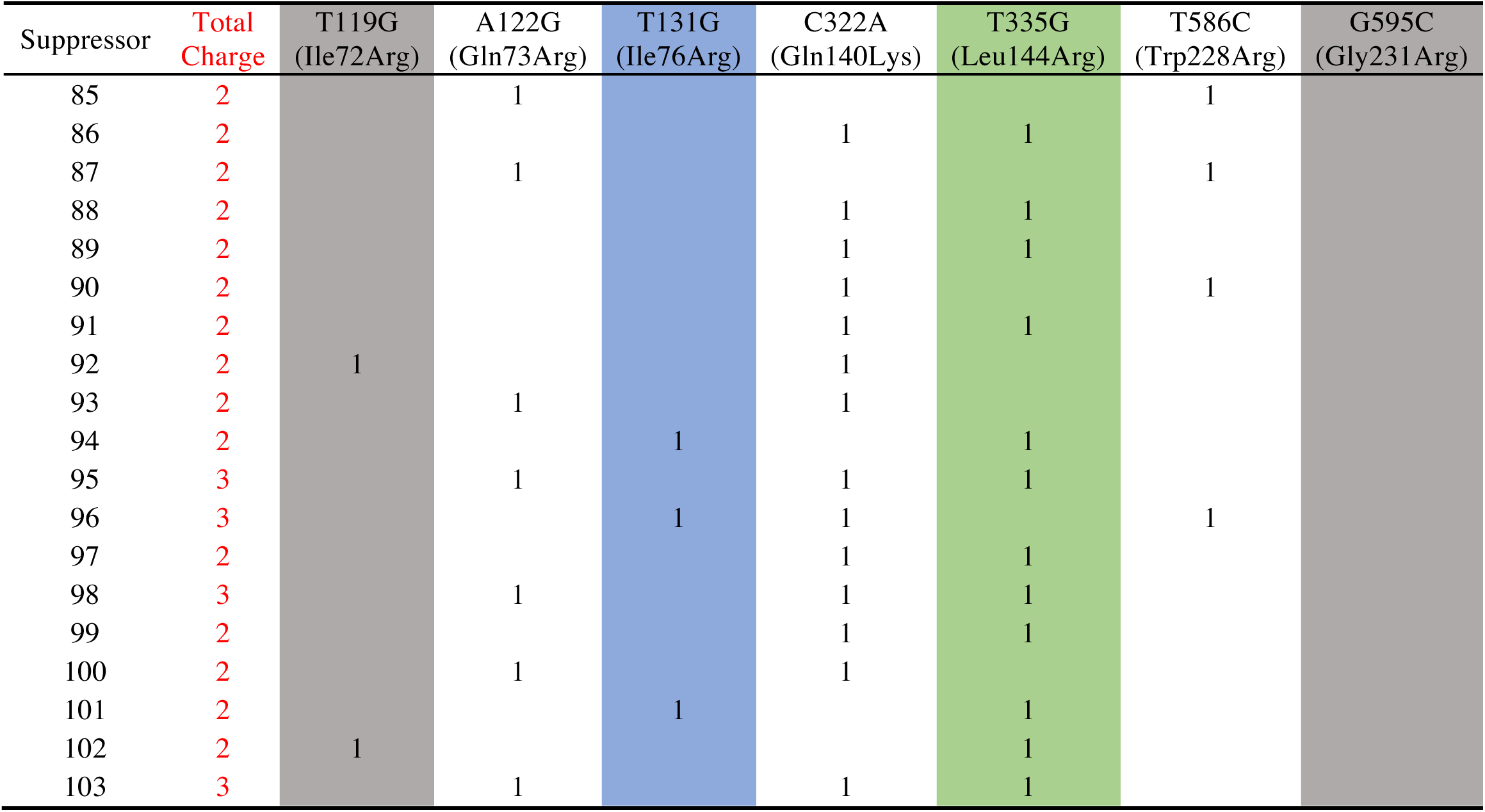
Amino acid substitutions in SpoIIIJ variants selected in *yhdL* depletion strain.

**Table S4.** Secreted and membrane-associated proteins in the σ^M^ regulon.

| Locus | Gene | Protein Localization | Number of predicted transmembrane segments |
| --- | --- | --- | --- |
| BSU00570 | *yabM* | cytoplasmic membrane | 14 |
| BSU00600 | *yabP* | forespore outer membrane | Unknown |
| BSU00610 | *yabQ* | forespore outer membrane | 6 |
| BSU00620 | *divIC* | peripheral membrane protein | Peripheral |
| BSU00690 | *ftsH* | cytoplasmic membrane | 2 |
| BSU02900 | *yceD* | peripheral membrane protein | Peripheral |
| BSU02920 | *yceF* | cytoplasmic membrane | 6 |
| BSU03240 | *ycgQ* | cytoplasmic membrane | 4 |
| BSU03250 | *ycgR* | cytoplasmic membrane | 8 |
| BSU04230 | *amj* | cytoplasmic membrane | 7 |
| BSU06380 | *yebC* | cytoplasmic membrane | 5 |
| BSU09500 | *yhdK* | cytoplasmic membrane | 3 |
| BSU09510 | *yhdL* | cytoplasmic membrane | 1 |
| BSU15220 | *murG* | peripheral membrane protein | Peripheral |
| BSU15240 | *divIB* | cytoplasmic membrane | 1 |
| BSU15250 | *ylxW* | Integral membrane protein | Unknown |
| BSU15260 | *ylxX* | cytoplasmic membrane | 1 |
| BSU15270 | *sbp* | cytoplasmic membrane | 3 |
| BSU15280 | *ftsA* | cytoplasmic membrane and cytoplasm | Peripheral |
| BSU15290 | *ftsZ* | cytoplasmic membrane and cytoplasm | Peripheral |
| BSU18190 | *yngC* | cytoplasmic membrane | 4 |
| BSU21920 | *ugtP* | peripheral membrane protein | Peripheral |
| BSU22320 | *ponA* | cytoplasmic membrane | 1 |
| BSU27160 | *yrhJ* | cytoplasmic membrane | Unknown |
| BSU27650 | *secDF* | cytoplasmic membrane | 12 |
| BSU27990 | *minD* | peripheral membrane protein | Peripheral |
| BSU28000 | *minC* | peripheral membrane protein | Peripheral |
| BSU28010 | *mreD* | cytoplasmic membrane | 5 |
| BSU28020 | *mreC* | cytoplasmic membrane | 1 |
| BSU28030 | *mreB* | peripheral membrane protein | Peripheral |
| BSU35650 | *tagU* | cytoplasmic membrane | 1 |
| BSU35840 | *tagT* | cytoplasmic membrane | 1 |
| BSU36530 | *bcrC* | cytoplasmic membrane | 4 |
| BSU38120 | *rodA* | cytoplasmic membrane | 11 |
| BSU38260 | *efeB* | Secreted protein | Secreted |
| BSU38270 | *efeO* | cytoplasmic membrane | Peripheral |
| BSU38280 | *efeU* | cytoplasmic membrane | 6 |
| BSU38510 | *dltB* | cytoplasmic membrane | 10 |

**Table S5.**
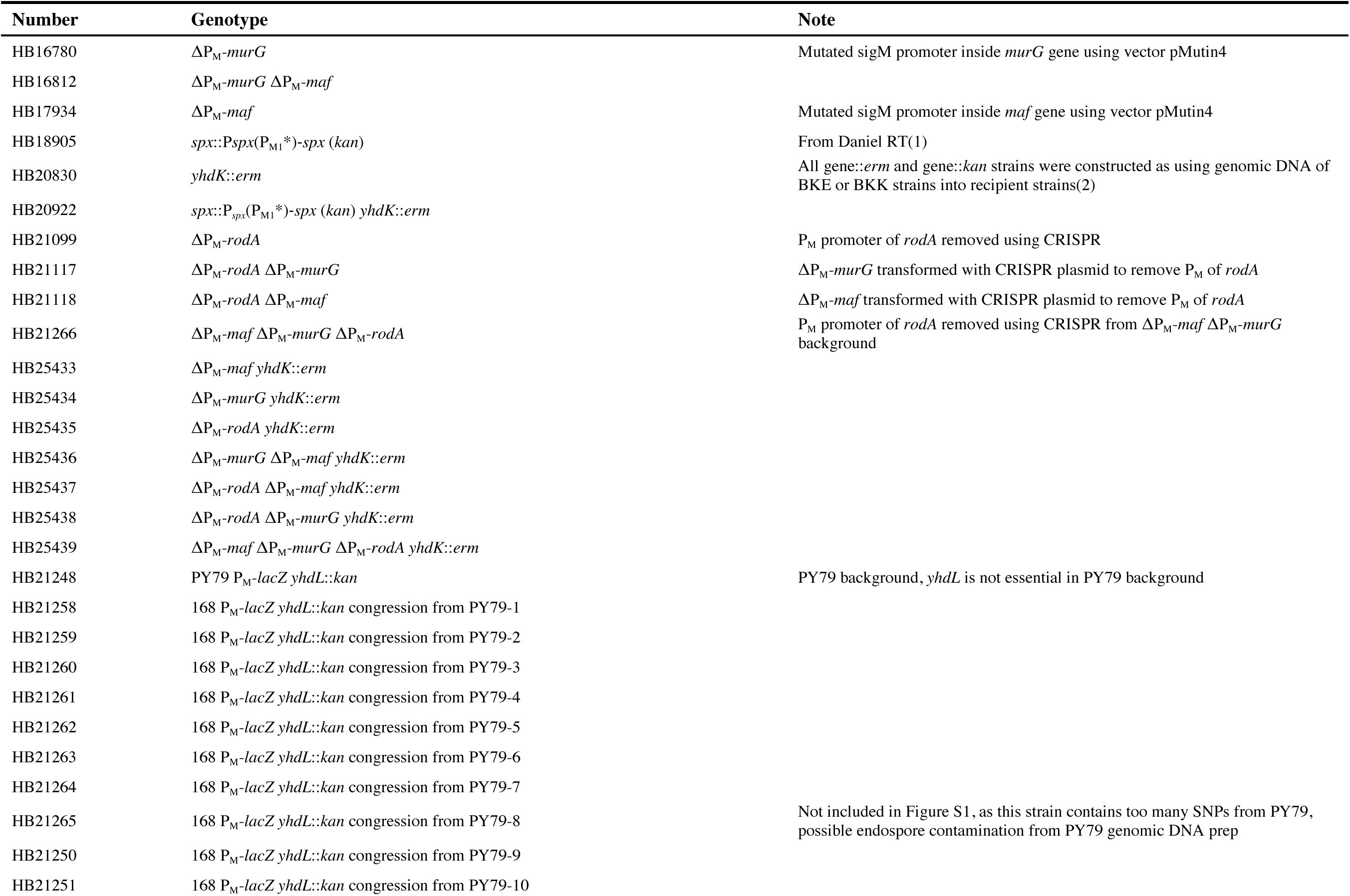
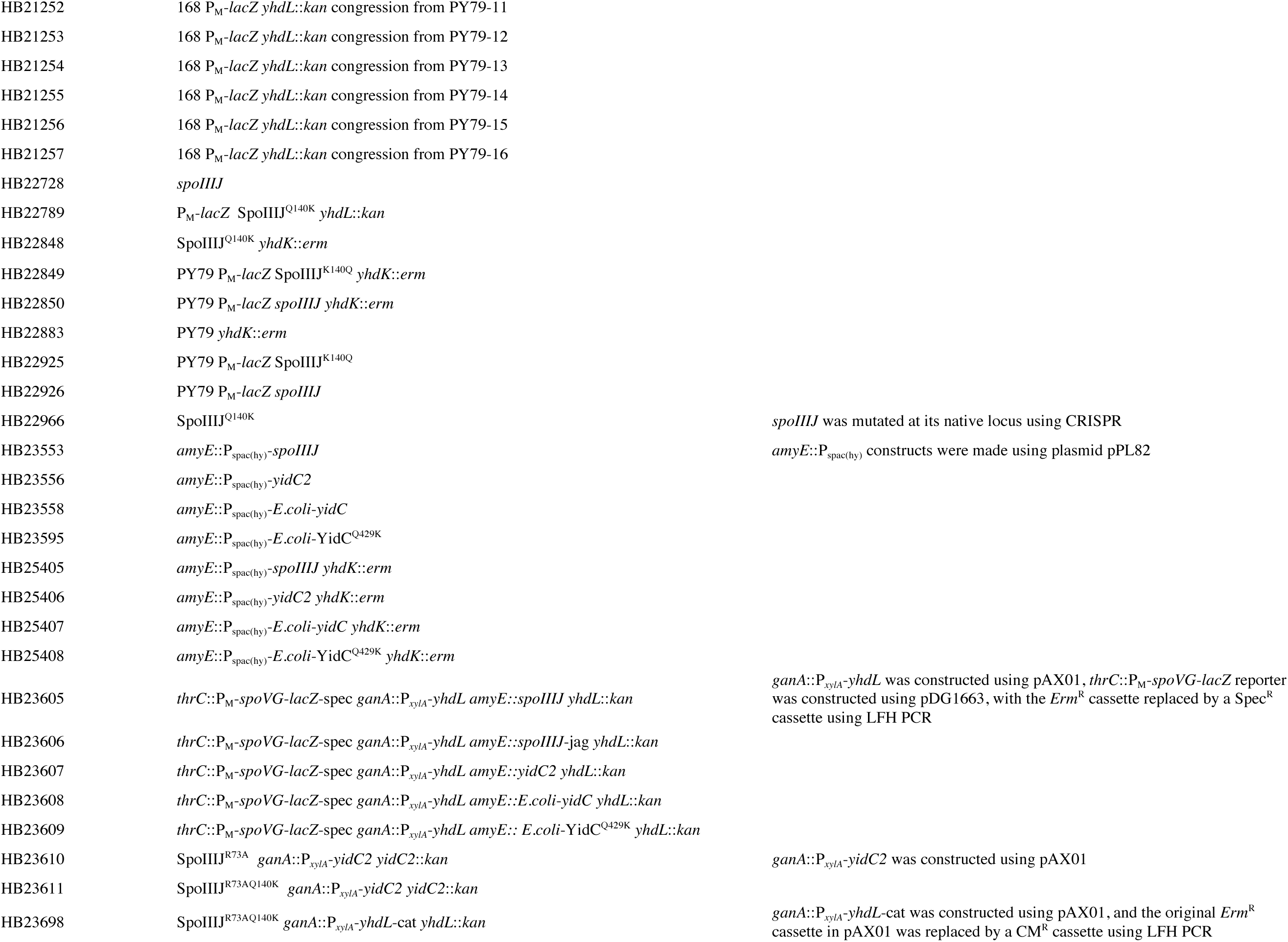
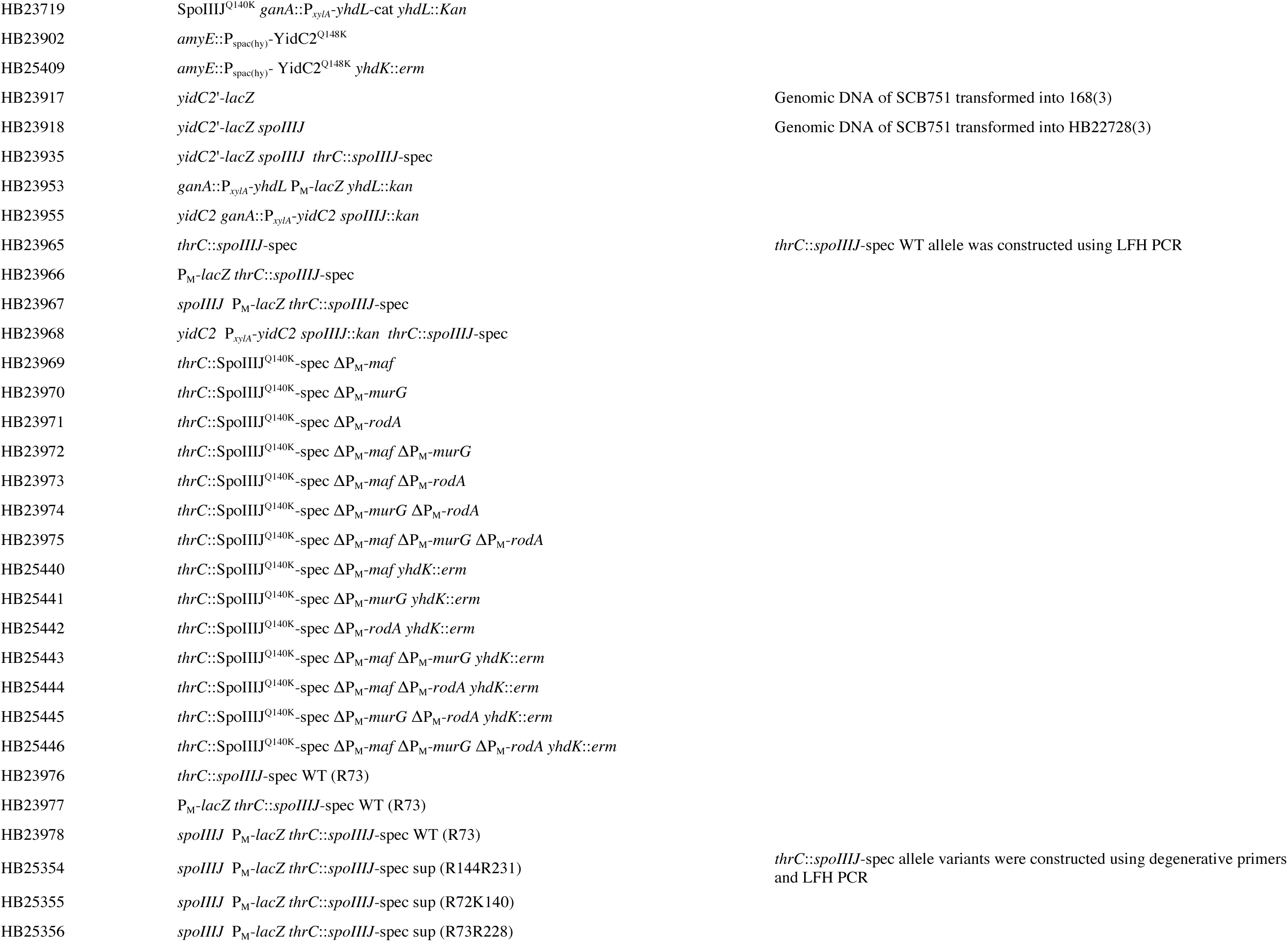
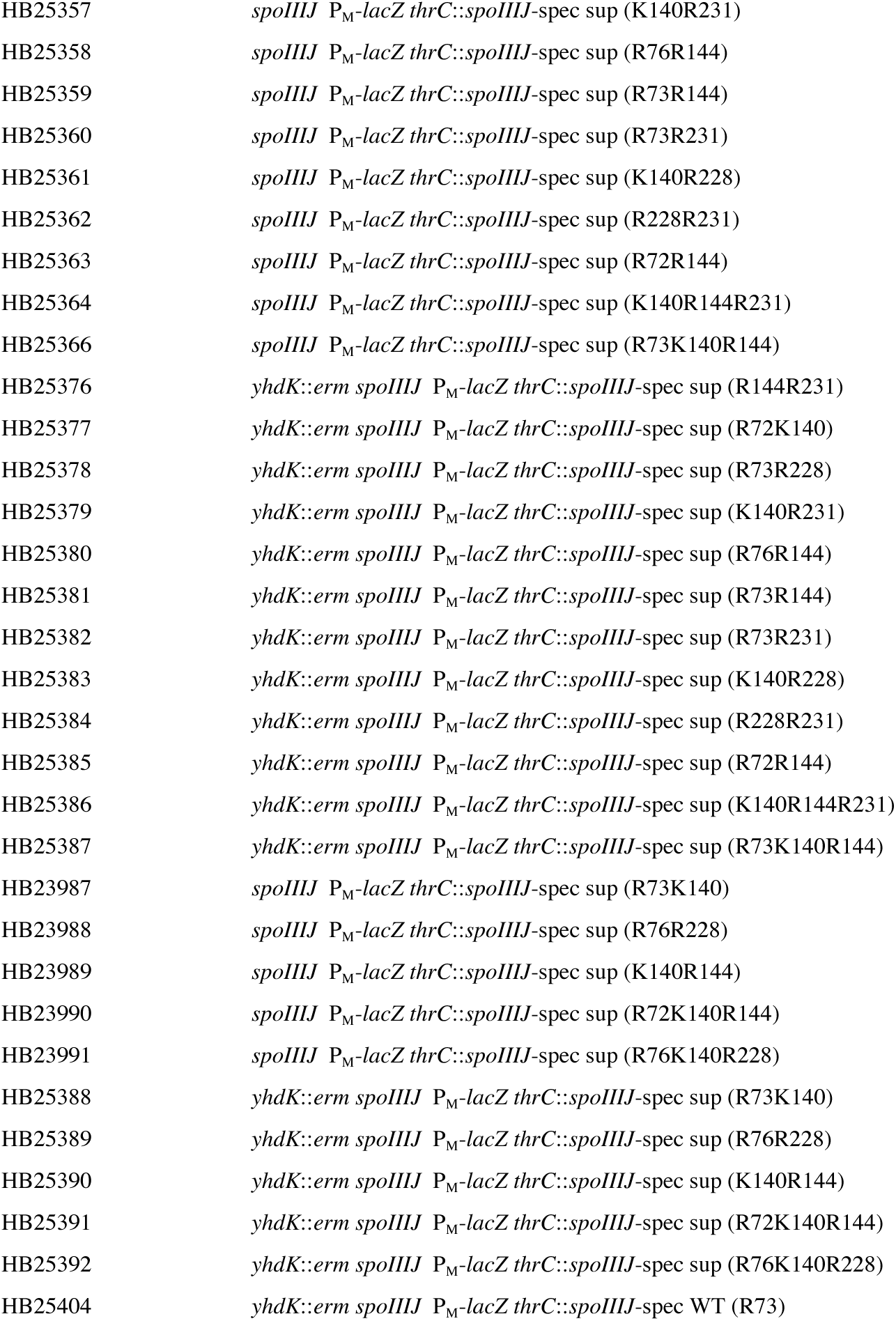
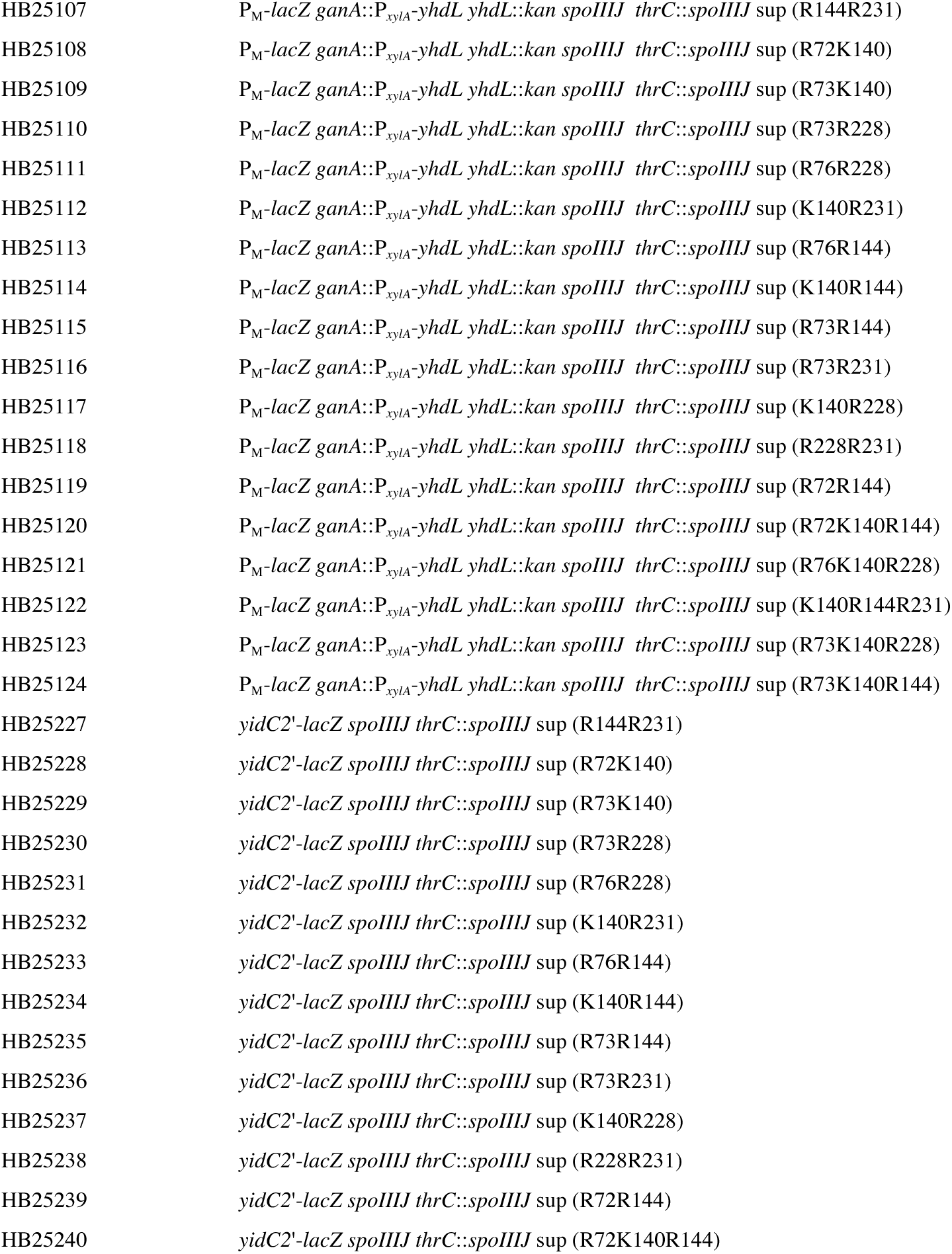
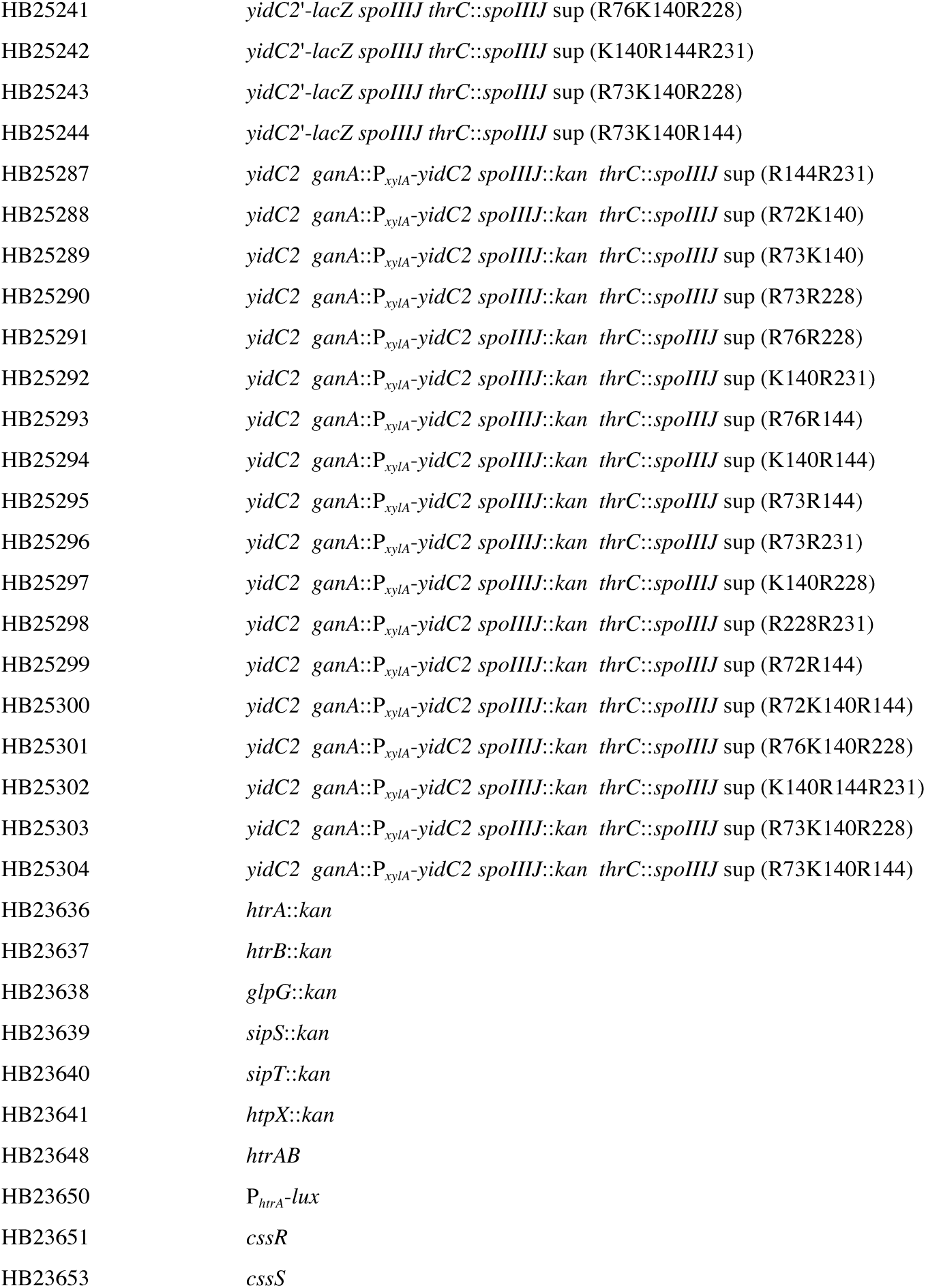
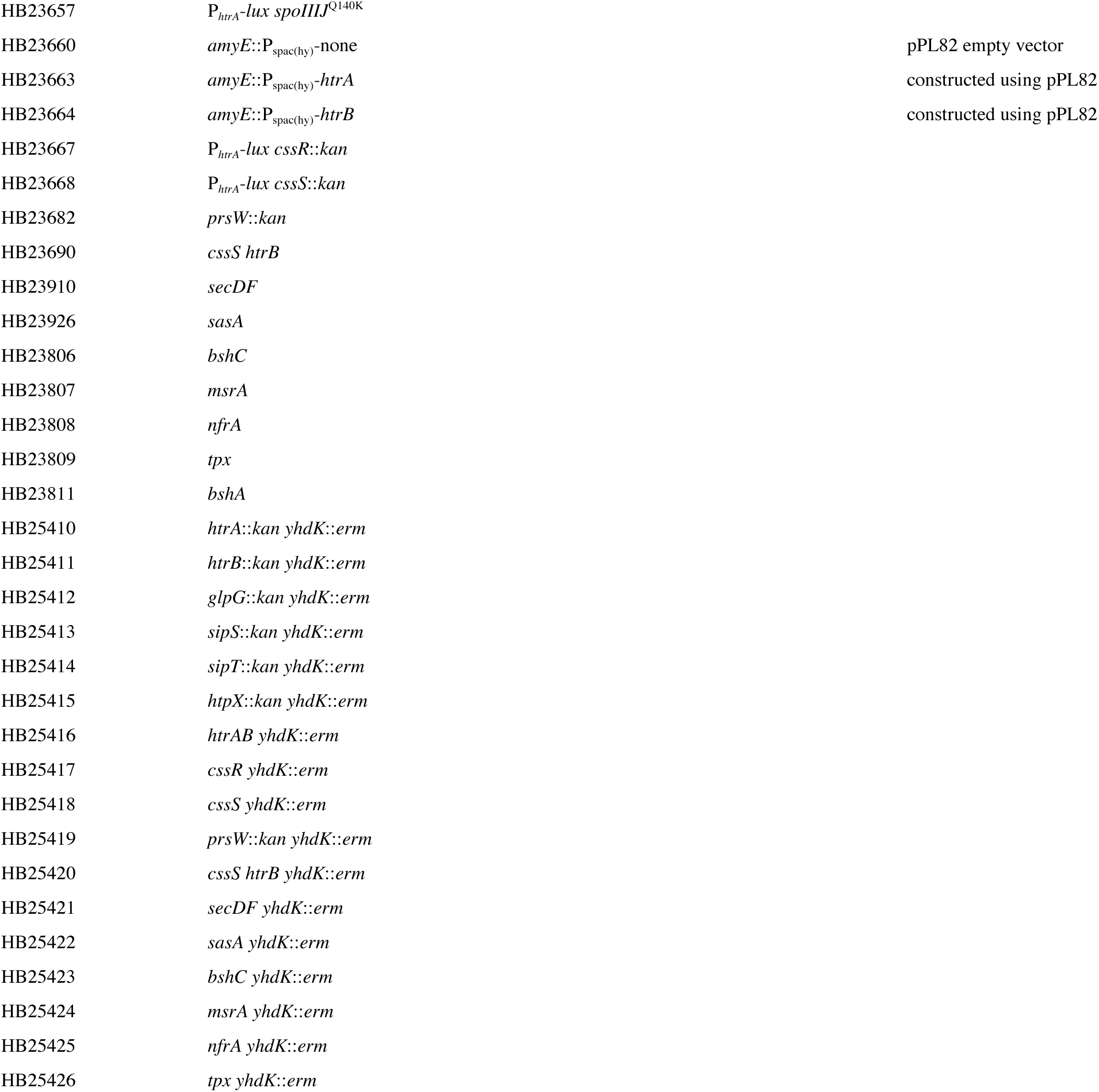
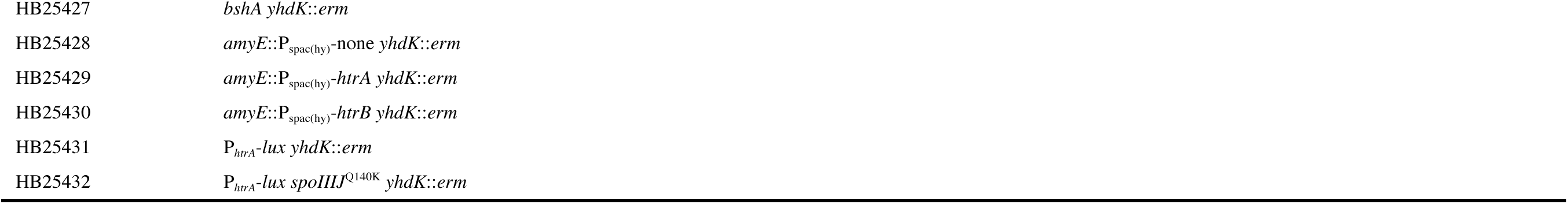
*Bacillus subtilis* strains used in this study.

**Table S6.**
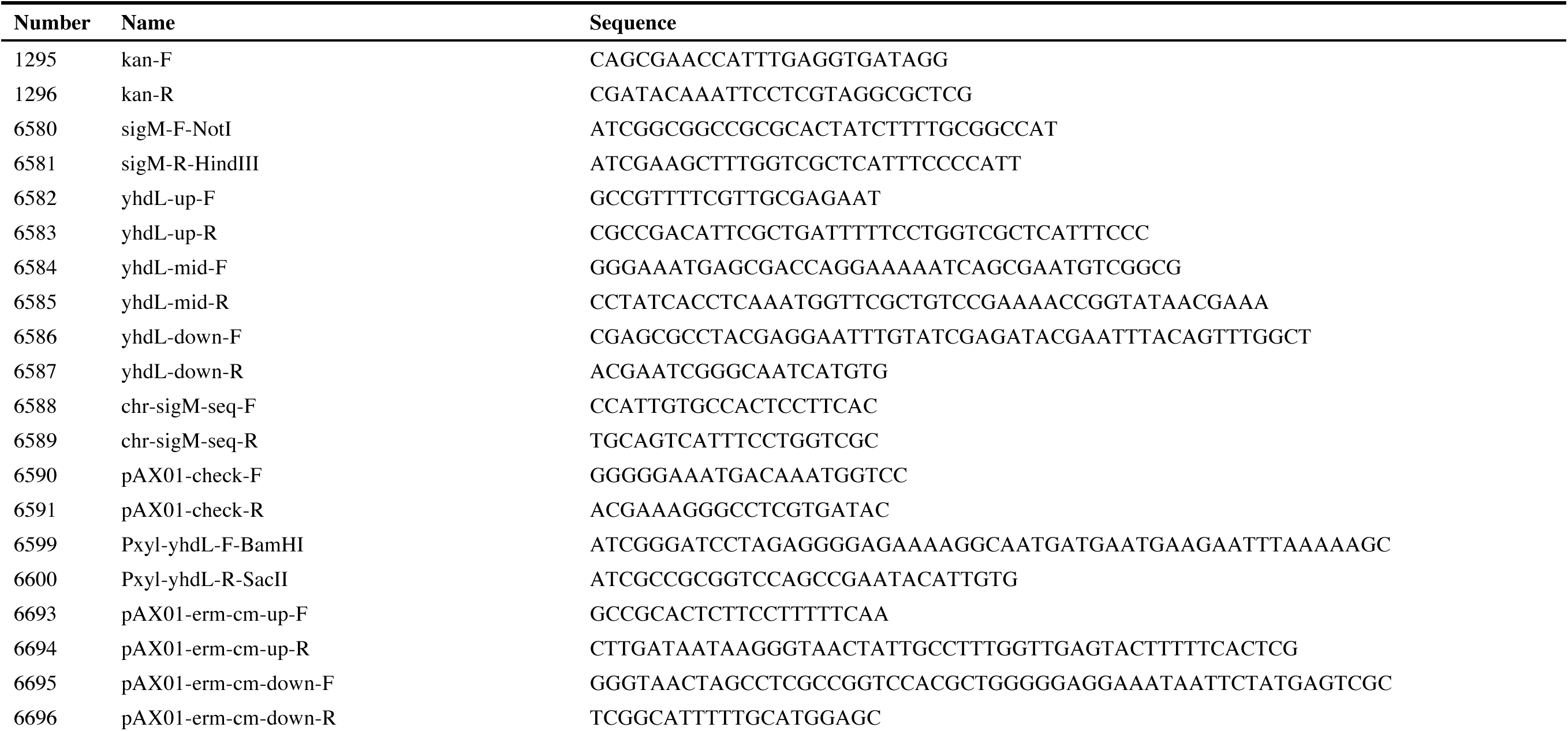
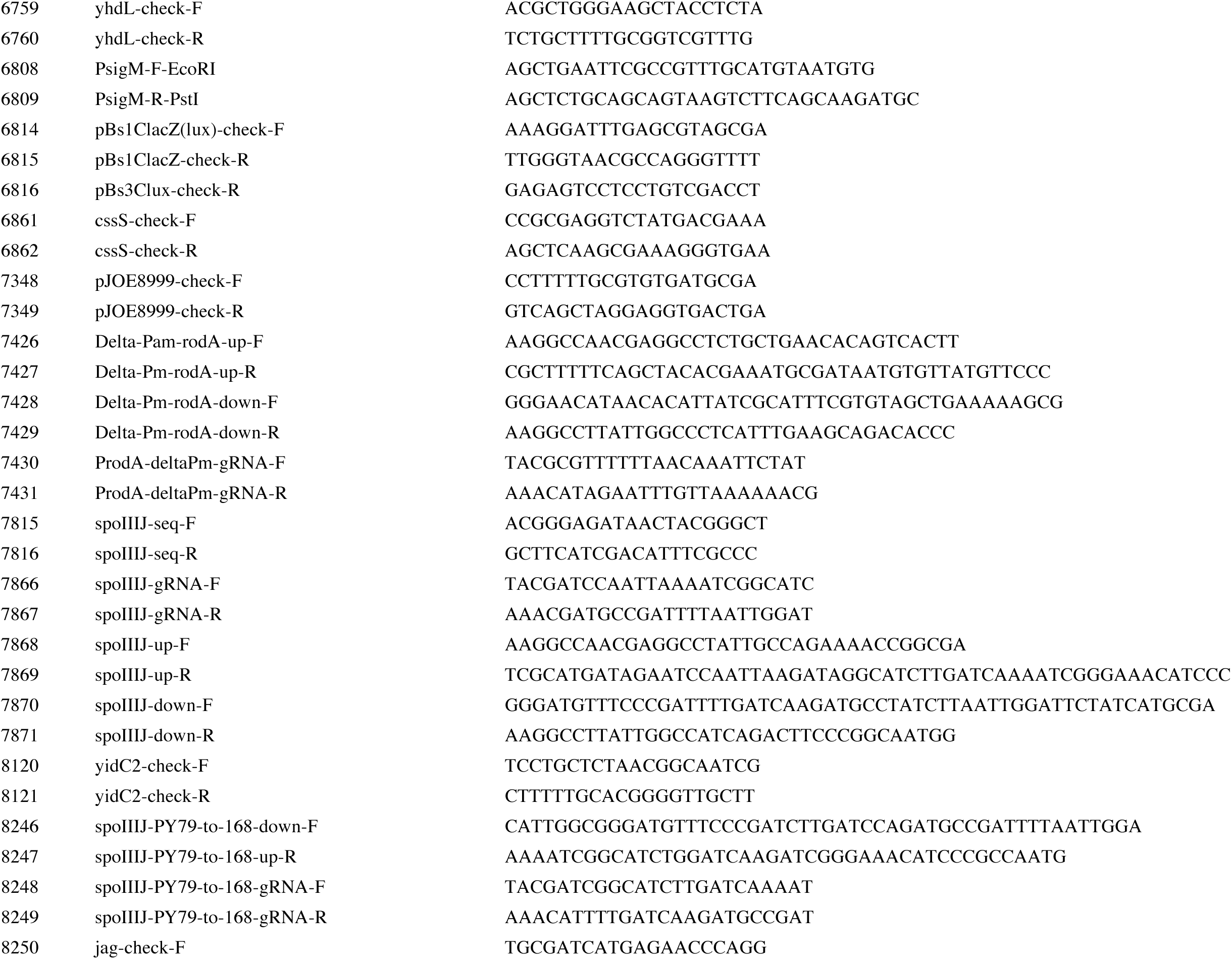
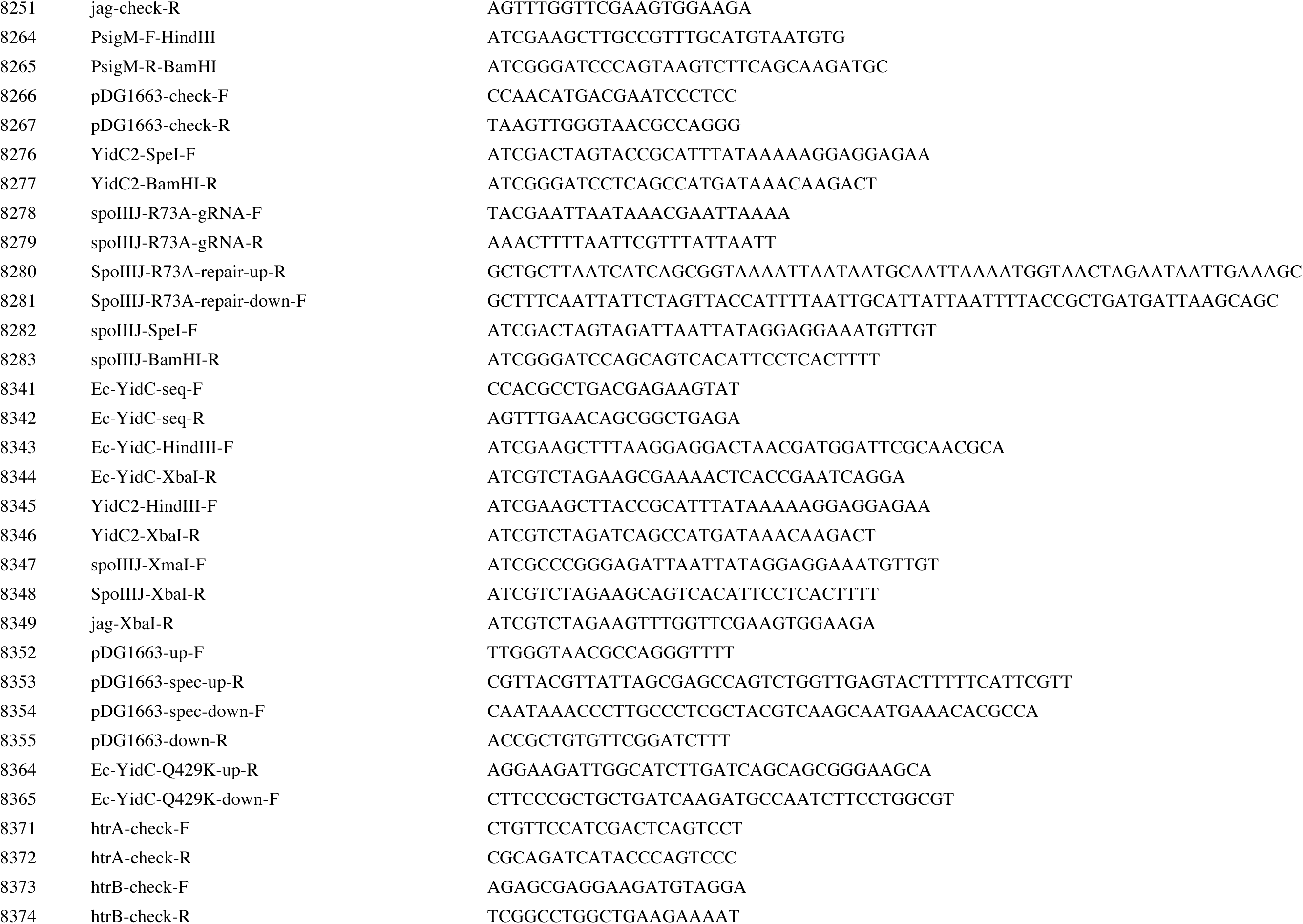
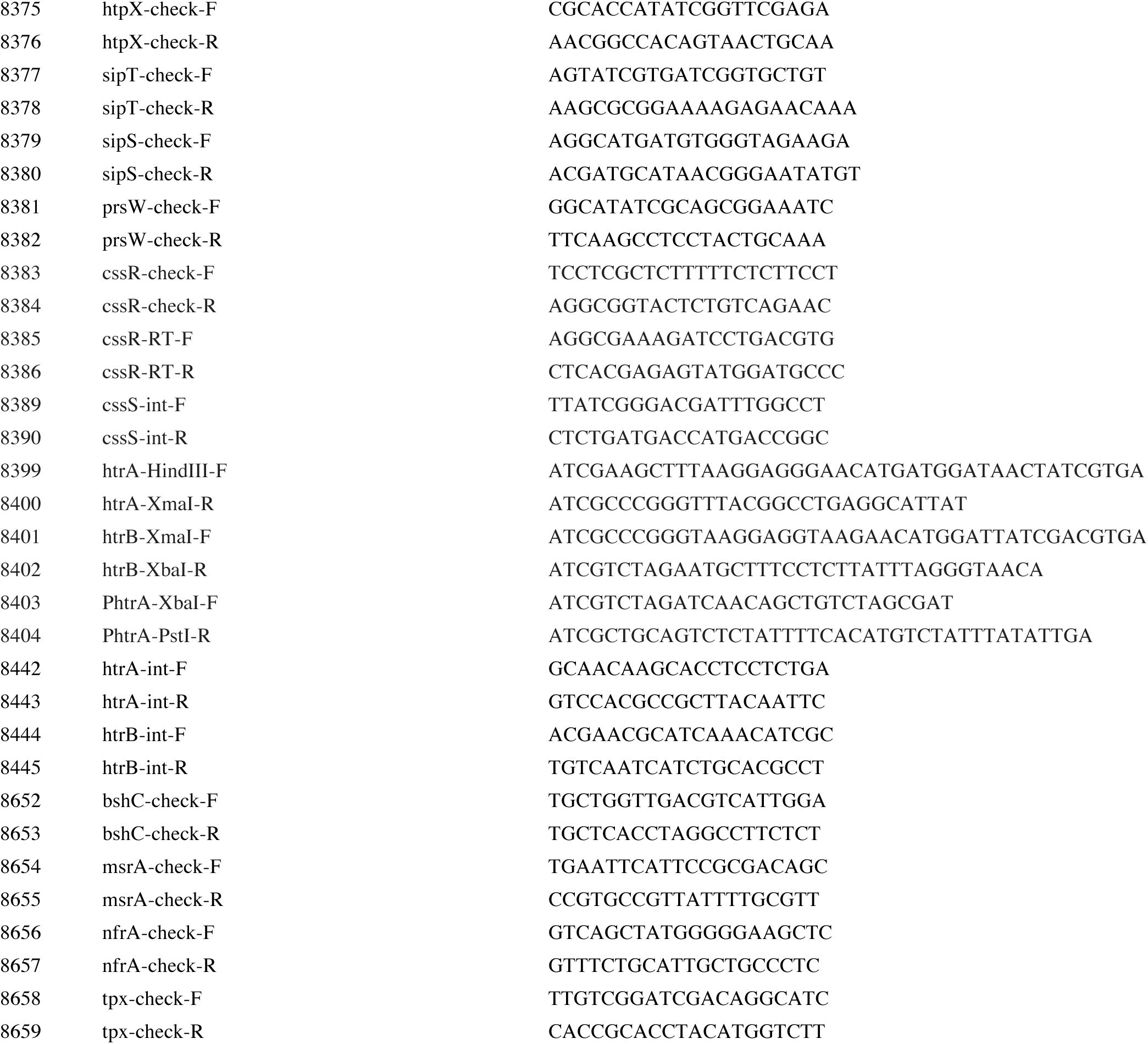
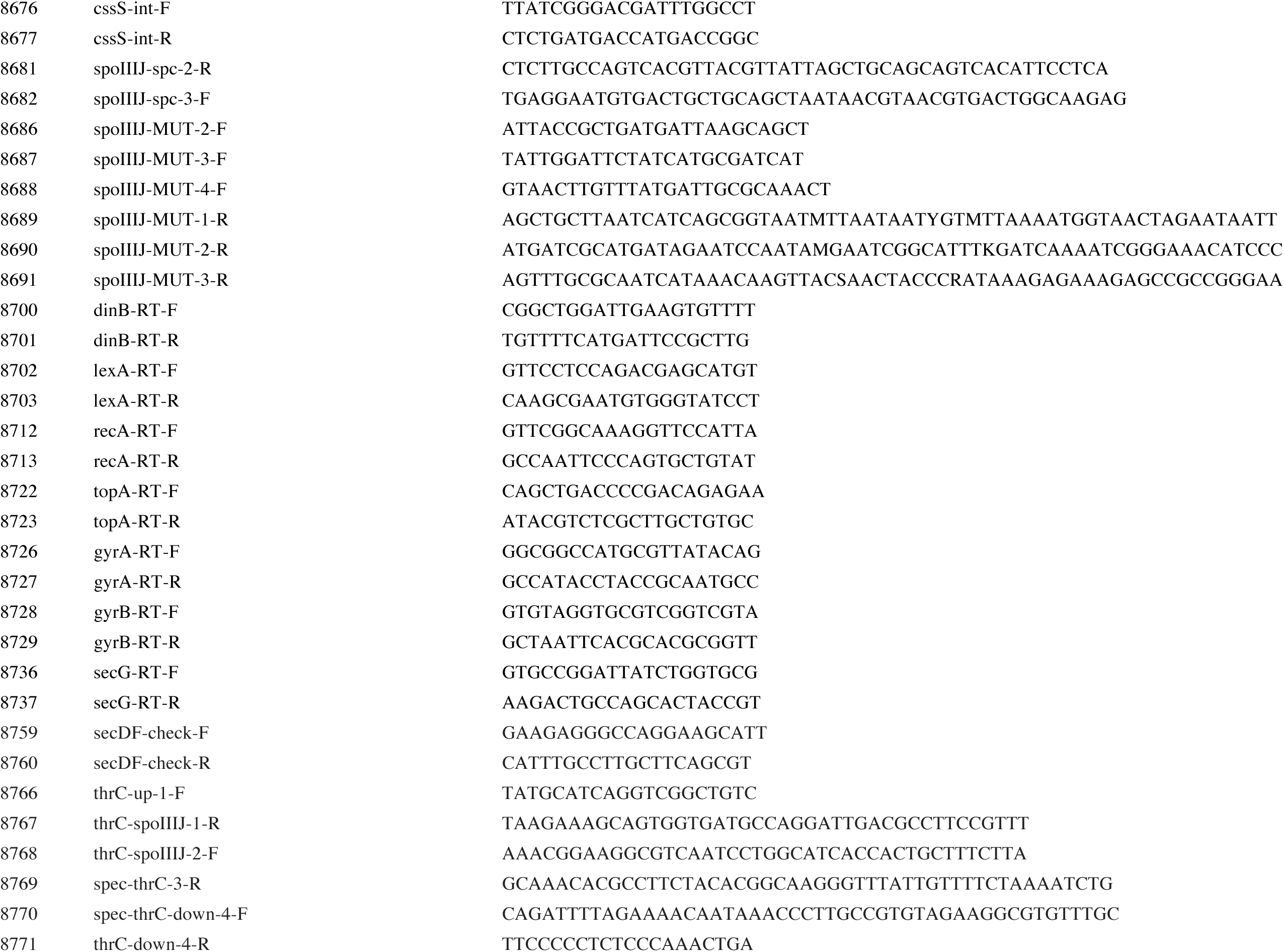
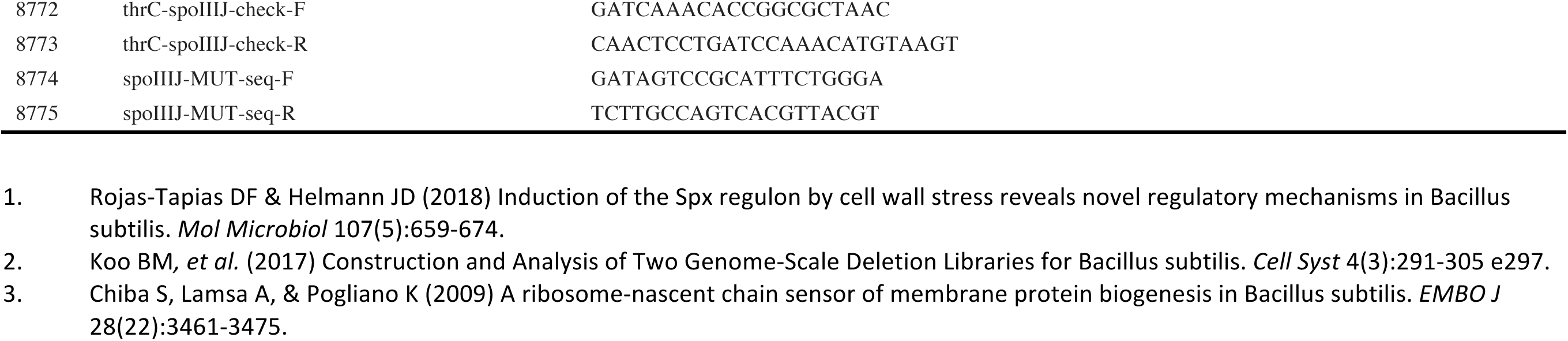
Primers used in this study.

## References

1. Helmann JD. *Bacillus subtilis* extracytoplasmic function (ECF) sigma factors and defense of the cell envelope. Curr Opin Microbiol. 2016;30:122–32. Epub 2016/02/24. doi: 10.1016/j.mib.2016.02.002. PubMed PMID: 26901131; PubMed Central PMCID: PMCPMC4821709.

2. Sineva E, Savkina M, Ades SE. Themes and variations in gene regulation by extracytoplasmic function (ECF) sigma factors. Curr Opin Microbiol. 2017;36:128–37. doi: 10.1016/j.mib.2017.05.004. PubMed PMID: 28575802; PubMed Central PMCID: PMCPMC5534382.

3. Eiamphungporn W, Helmann JD. The *Bacillus subtilis* sigma(M) regulon and its contribution to cell envelope stress responses. Mol Microbiol. 2008;67(4):830–48. Epub 2008/01/09. doi: 10.1111/j.1365-2958.2007.06090.x. PubMed PMID: 18179421; PubMed Central PMCID: PMCPMC3025603.

4. Horsburgh MJ, Moir A. Sigma M, an ECF RNA polymerase sigma factor of *Bacillus subtilis* 168, is essential for growth and survival in high concentrations of salt. Mol Microbiol. 1999;32(1):41–50. Epub 1999/04/27. PubMed PMID: 10216858.

5. Luo Y, Helmann JD. Analysis of the role of *Bacillus subtilis* sigma(M) in beta-lactam resistance reveals an essential role for c-di-AMP in peptidoglycan homeostasis. Mol Microbiol. 2012;83(3):623–39. Epub 2012/01/04. doi: 10.1111/j.1365-2958.2011.07953.x. PubMed PMID: 22211522; PubMed Central PMCID: PMCPMC3306796.

6. Meeske AJ, Riley EP, Robins WP, Uehara T, Mekalanos JJ, Kahne D, et al. SEDS proteins are a widespread family of bacterial cell wall polymerases. Nature. 2016;537(7622):634–8. Epub 2016/08/16. doi: 10.1038/nature19331. PubMed PMID: 27525505; PubMed Central PMCID: PMCPMC5161649.

7. Asai K. Anti-sigma factor-mediated cell surface stress responses in *Bacillus subtilis*. Genes Genet Syst. 2018;92(5):223–34. Epub 2018/01/19. doi: 10.1266/ggs.17-00046. PubMed PMID: 29343670.

8. Woods EC, McBride SM. Regulation of antimicrobial resistance by extracytoplasmic function (ECF) sigma factors. Microbes Infect. 2017;19(4-5):238–48. doi: 10.1016/j.micinf.2017.01.007. PubMed PMID: 28153747; PubMed Central PMCID: PMCPMC5403605.

9. Yoshimura M, Asai K, Sadaie Y, Yoshikawa H. Interaction of *Bacillus subtilis* extracytoplasmic function (ECF) sigma factors with the N-terminal regions of their potential anti-sigma factors. Microbiology. 2004;150(Pt 3):591–9. doi: 10.1099/mic.0.26712-0. PubMed PMID: 14993308.

10. Zhao H, Roistacher DM, Helmann JD. Deciphering the essentiality and function of the anti-sigma(M) factors in *Bacillus subtilis*. Mol Microbiol. 2019. doi: 10.1111/mmi.14216. PubMed PMID: 30715747.

11. Murakami T, Haga K, Takeuchi M, Sato T. Analysis of the *Bacillus subtilis spoIIIJ* gene and its Paralogue gene, *yqjG*. J Bacteriol. 2002;184(7):1998–2004. doi: 10.1128/jb.184.7.1998-2004.2002. PubMed PMID: 11889108; PubMed Central PMCID: PMCPMC134917.

12. Tjalsma H, Bron S, van Dijl JM. Complementary impact of paralogous Oxa1-like proteins of *Bacillus subtilis* on post-translocational stages in protein secretion. J Biol Chem. 2003;278(18):15622–32. doi: 10.1074/jbc.M301205200. PubMed PMID: 12586834.

13. Zeigler DR, Pragai Z, Rodriguez S, Chevreux B, Muffler A, Albert T, et al. The origins of 168, W23, and other Bacillus subtilis legacy strains. J Bacteriol. 2008;190(21):6983–95. Epub 2008/08/30. doi: 10.1128/JB.00722-08. PubMed PMID: 18723616; PubMed Central PMCID: PMCPMC2580678.

14. Schroeder JW, Simmons LA. Complete Genome Sequence of *Bacillus subtilis* Strain PY79. Genome Announc. 2013;1(6). Epub 2013/12/21. doi: 10.1128/genomeA.01085-13. PubMed PMID: 24356846; PubMed Central PMCID: PMCPMC3868870.

15. Dubnau D, Cirigliano C. Fate of transforming DNA following uptake by competent *Bacillus subtilis*. Formation and properties of products isolated from transformed cells which are derived entirely from donor DNA. J Mol Biol. 1972;64(1):9–29. Epub 1972/02/28. PubMed PMID: 4622632.

16. Kumazaki K, Chiba S, Takemoto M, Furukawa A, Nishiyama K, Sugano Y, et al. Structural basis of Sec-independent membrane protein insertion by YidC. Nature. 2014;509(7501):516–20. Epub 2014/04/18. doi: 10.1038/nature13167. PubMed PMID: 24739968.

17. Tsirigotaki A, De Geyter J, Sostaric N, Economou A, Karamanou S. Protein export through the bacterial Sec pathway. Nat Rev Microbiol. 2017;15(1):21–36. Epub 2016/11/29. doi: 10.1038/nrmicro.2016.161. PubMed PMID: 27890920.

18. Hennon SW, Soman R, Zhu L, Dalbey RE. YidC/Alb3/Oxa1 Family of Insertases. J Biol Chem. 2015;290(24):14866–74. Epub 2015/05/08. doi: 10.1074/jbc.R115.638171. PubMed PMID: 25947384; PubMed Central PMCID: PMCPMC4463434.

19. Saller MJ, Fusetti F, Driessen AJ. *Bacillus subtilis* SpoIIIJ and YqjG function in membrane protein biogenesis. J Bacteriol. 2009;191(21):6749–57. Epub 2009/09/01. doi: 10.1128/JB.00853-09. PubMed PMID: 19717609; PubMed Central PMCID: PMCPMC2795313.

20. Errington J, Appleby L, Daniel RA, Goodfellow H, Partridge SR, Yudkin MD. Structure and function of the *spoIIIJ* gene of *Bacillus subtilis*: a vegetatively expressed gene that is essential for sigma G activity at an intermediate stage of sporulation. J Gen Microbiol. 1992;138(12):2609–18. Epub 1992/12/01. doi: 10.1099/00221287-138-12-2609. PubMed PMID: 1487728.

21. Chiba S, Ito K. MifM monitors total YidC activities of *Bacillus subtilis*, including that of YidC2, the target of regulation. J Bacteriol. 2015;197(1):99–107. Epub 2014/10/15. doi: 10.1128/JB.02074-14. PubMed PMID: 25313395; PubMed Central PMCID: PMCPMC4288694.

22. Corte L, Valente F, Serrano M, Gomes CM, Moran CP, Jr., Henriques AO. A conserved cysteine residue of *Bacillus subtilis* SpoIIIJ is important for endospore development. PLoS One. 2014;9(8):e99811. Epub 2014/08/19. doi: 10.1371/journal.pone.0099811. PubMed PMID: 25133632; PubMed Central PMCID: PMCPMC4136701.

23. Saller MJ, Otto A, Berrelkamp-Lahpor GA, Becher D, Hecker M, Driessen AJ. *Bacillus subtilis* YqjG is required for genetic competence development. Proteomics. 2011;11(2):270–82. Epub 2011/01/05. doi: 10.1002/pmic.201000435. PubMed PMID: 21204254.

24. Quisel JD, Burkholder WF, Grossman AD. In vivo effects of sporulation kinases on mutant Spo0A proteins in *Bacillus subtilis*. J Bacteriol. 2001;183(22):6573–8. doi: 10.1128/JB.183.22.6573-6578.2001. PubMed PMID: 11673427; PubMed Central PMCID: PMCPMC95488.

25. Shimokawa-Chiba N, Kumazaki K, Tsukazaki T, Nureki O, Ito K, Chiba S. Hydrophilic microenvironment required for the channel-independent insertase function of YidC protein. Proc Natl Acad Sci U S A. 2015;112(16):5063–8. Epub 2015/04/10. doi: 10.1073/pnas.1423817112. PubMed PMID: 25855636; PubMed Central PMCID: PMCPMC4413333.

26. Chiba S, Ito K. Multisite ribosomal stalling: a unique mode of regulatory nascent chain action revealed for MifM. Mol Cell. 2012;47(6):863–72. Epub 2012/08/07. doi: 10.1016/j.molcel.2012.06.034. PubMed PMID: 22864117.

27. Kuhn A, Kiefer D. Membrane protein insertase YidC in bacteria and archaea. Mol Microbiol. 2017;103(4):590–4. Epub 2016/11/24. doi: 10.1111/mmi.13586. PubMed PMID: 27879020.

28. Hyyrylainen HL, Bolhuis A, Darmon E, Muukkonen L, Koski P, Vitikainen M, et al. A novel two-component regulatory system in *Bacillus subtilis* for the survival of severe secretion stress. Mol Microbiol. 2001;41(5):1159–72. PubMed PMID: 11555295.

29. Darmon E, Noone D, Masson A, Bron S, Kuipers OP, Devine KM, et al. A novel class of heat and secretion stress-responsive genes is controlled by the autoregulated CssRS two-component system of *Bacillus subtilis*. J Bacteriol. 2002;184(20):5661–71. doi: 10.1128/jb.184.20.5661-5671.2002. PubMed PMID: 12270824; PubMed Central PMCID: PMCPMC139597.

30. Podgornaia AI, Laub MT. Determinants of specificity in two-component signal transduction. Curr Opin Microbiol. 2013;16(2):156–62. Epub 2013/01/29. doi: 10.1016/j.mib.2013.01.004. PubMed PMID: 23352354.

31. Takahashi N, Gruber CC, Yang JH, Liu X, Braff D, Yashaswini CN, et al. Lethality of MalE-LacZ hybrid protein shares mechanistic attributes with oxidative component of antibiotic lethality. Proc Natl Acad Sci U S A. 2017. Epub 2017/08/11. doi: 10.1073/pnas.1707466114. PubMed PMID: 28794281; PubMed Central PMCID: PMCPMC5576823.

32. Kohanski MA, Dwyer DJ, Wierzbowski J, Cottarel G, Collins JJ. Mistranslation of membrane proteins and two-component system activation trigger antibiotic-mediated cell death. Cell. 2008;135(4):679–90. Epub 2008/11/18. doi: 10.1016/j.cell.2008.09.038. PubMed PMID: 19013277; PubMed Central PMCID: PMCPMC2684502.

33. Zhu B, Stulke J. SubtiWiki in 2018: from genes and proteins to functional network annotation of the model organism Bacillus subtilis. Nucleic Acids Res. 2018;46(D1):D743–D8. Epub 2018/05/23. doi: 10.1093/nar/gkx908. PubMed PMID: 29788229; PubMed Central PMCID: PMCPMC5753275.

34. Price CE, Driessen AJ. YidC is involved in the biogenesis of anaerobic respiratory complexes in the inner membrane of *Escherichia coli*. J Biol Chem. 2008;283(40):26921–7. doi: 10.1074/jbc.M804490200. PubMed PMID: 18635537.

35. Dalbey RE, Kuhn A, Zhu L, Kiefer D. The membrane insertase YidC. Biochim Biophys Acta. 2014;1843(8):1489–96. doi: 10.1016/j.bbamcr.2013.12.022. PubMed PMID: 24418623.

36. Chiba S, Lamsa A, Pogliano K. A ribosome-nascent chain sensor of membrane protein biogenesis in *Bacillus subtilis*. EMBO J. 2009;28(22):3461–75. Epub 2009/09/26. doi: 10.1038/emboj.2009.280. PubMed PMID: 19779460; PubMed Central PMCID: PMCPMC2782093.

37. Chen Y, Soman R, Shanmugam SK, Kuhn A, Dalbey RE. The role of the strictly conserved positively charged residue differs among the Gram-positive, Gram-negative, and chloroplast YidC homologs. J Biol Chem. 2014;289(51):35656–67. Epub 2014/11/02. doi: 10.1074/jbc.M114.595082. PubMed PMID: 25359772; PubMed Central PMCID: PMCPMC4271247.

38. Luo Y, Asai K, Sadaie Y, Helmann JD. Transcriptomic and phenotypic characterization of a *Bacillus subtilis* strain without extracytoplasmic function sigma factors. J Bacteriol. 2010;192(21):5736–45. Epub 2010/09/08. doi: 10.1128/JB.00826-10. PubMed PMID: 20817771; PubMed Central PMCID: PMCPMC2953670.

39. Zhao H, Sun Y, Peters JM, Gross CA, Garner EC, Helmann JD. Depletion of Undecaprenyl Pyrophosphate Phosphatases Disrupts Cell Envelope Biogenesis in *Bacillus subtilis*. J Bacteriol. 2016;198(21):2925–35. doi: 10.1128/JB.00507-16. PubMed PMID: 27528508; PubMed Central PMCID: PMCPMC5055597.

40. Hartl B, Wehrl W, Wiegert T, Homuth G, Schumann W. Development of a new integration site within the *Bacillus subtilis* chromosome and construction of compatible expression cassettes. J Bacteriol. 2001;183(8):2696–9. Epub 2001/03/29. doi: 10.1128/JB.183.8.2696-2699.2001. PubMed PMID: 11274134; PubMed Central PMCID: PMCPMC95191.

41. Koo BM, Kritikos G, Farelli JD, Todor H, Tong K, Kimsey H, et al. Construction and Analysis of Two Genome-Scale Deletion Libraries for *Bacillus subtilis*. Cell Syst. 2017;4(3):291–305 e7. Epub 2017/02/13. doi: 10.1016/j.cels.2016.12.013. PubMed PMID: 28189581; PubMed Central PMCID: PMCPMC5400513.

42. Rojas-Tapias DF, Helmann JD. Induction of the Spx regulon by cell wall stress reveals novel regulatory mechanisms in *Bacillus subtilis*. Mol Microbiol. 2018;107(5):659–74. Epub 2017/12/23. doi: 10.1111/mmi.13906. PubMed PMID: 29271514; PubMed Central PMCID: PMCPMC5820111.

43. Altenbuchner J. Editing of the *Bacillus subtilis* Genome by the CRISPR-Cas9 System. Appl Environ Microbiol. 2016;82(17):5421–7. Epub 2016/06/28. doi: 10.1128/AEM.01453-16. PubMed PMID: 27342565; PubMed Central PMCID: PMCPMC4988203.

44. Moszer I, Rocha EP, Danchin A. Codon usage and lateral gene transfer in *Bacillus subtilis*. Curr Opin Microbiol. 1999;2(5):524–8. Epub 1999/10/06. PubMed PMID: 10508724.

45. Schindelin J, Arganda-Carreras I, Frise E, Kaynig V, Longair M, Pietzsch T, et al. Fiji: an open-source platform for biological-image analysis. Nat Methods. 2012;9(7):676–82. Epub 2012/06/30. doi: 10.1038/nmeth.2019. PubMed PMID: 22743772; PubMed Central PMCID: PMCPMC3855844.

46. Zhao H, Roistacher DM, Helmann JD. Aspartate deficiency limits peptidoglycan synthesis and sensitizes cells to antibiotics targeting cell wall synthesis in *Bacillus subtilis*. Mol Microbiol. 2018;109(6):826–44. Epub 2018/07/12. doi: 10.1111/mmi.14078. PubMed PMID: 29995990; PubMed Central PMCID: PMCPMC6185803.

47. Radeck J, Kraft K, Bartels J, Cikovic T, Durr F, Emenegger J, et al. The *Bacillus* BioBrick Box: generation and evaluation of essential genetic building blocks for standardized work with Bacillus subtilis. J Biol Eng. 2013;7(1):29. Epub 2013/12/04. doi: 10.1186/1754-1611-7-29. PubMed PMID: 24295448; PubMed Central PMCID: PMCPMC4177231.

48. Pettersen EF, Goddard TD, Huang CC, Couch GS, Greenblatt DM, Meng EC, et al. UCSF Chimera--a visualization system for exploratory research and analysis. J Comput Chem. 2004;25(13):1605–12. Epub 2004/07/21. doi: 10.1002/jcc.20084. PubMed PMID: 15264254.

